# Identification of residue pairing in interacting β-strands from a predicted residue contact map

**DOI:** 10.1101/214643

**Authors:** Wenzhi Mao, Tong Wang, Wenxuan Zhang, Haipeng Gong

## Abstract

Despite the rapid progress of protein residue contact prediction, predicted residue contact maps frequently contain many errors. However, information of residue pairing in β strands could be extracted from a noisy contact map, due to the presence of characteristic contact patterns in β-β interactions. This information may benefit the tertiary structure prediction of mainly β proteins. In this work, we introduce a novel ridge-detection-based β-β contact predictor, RDb_2_C, to identify residue pairing in β strands from any predicted residue contact map. The algorithm adopts ridge detection, a well-developed technique in computer image processing, to capture consecutive residue contacts, and then utilizes a novel multi-stage random forest framework to integrate the ridge information and additional features for prediction. Starting from the predicted contact map of CCMpred, RDb_2_C remarkably outperforms all state-of-the-art methods on two conventional test sets of β proteins (BetaSheet916 and BetaSheet1452), and achieves F1-scores of ~62% and ~76% at the residue level and strand level, respectively. Taking the prediction of the more advanced RaptorX-Contact as input, RDb_2_C achieves impressively higher performance, with F1-scores reaching ~76% and ~86% at the residue level and strand level, respectively. According to our tests on 61 mainly β proteins, improvement in the β-β contact prediction can further ameliorate the structural prediction.

Availability: All source data and codes are available at http://166.111.152.91/Downloads.html or at the GitHub address of https://github.com/wzmao/RDb2C.

**Author summary:** Due to the topological complexity, mainly β proteins are challenging targets in protein structure prediction. Knowledge of the pairing between β strands, especially the residue pairing pattern, can greatly facilitate the tertiary structure prediction of mainly β proteins. In this work, we developed a novel algorithm to identify the residue pairing in β strands from a predicted residue contact map. This method adopts the ridge detection technique to capture the characteristic pattern of β-β interactions from the map and then utilizes a multi-stage random forest framework to predict β-β contacts at the residue level. According to our tests, our method could effectively improve the prediction of β-β contacts even from a highly noisy contact map. Moreover, the refined β-β contact information could effectively improve the structural modeling of mainly β proteins.

## Introduction

Since Anfinsen's dogma [1] was firstly introduced, prediction of the tertiary structures of proteins has become the Holy Grail in structural bioinformatics. Although practical tertiary structure prediction generally requires intensive sampling in the conformational space, the computational consumption could be greatly alleviated with the knowledge of residue contacts in the native conformation. Consequently, protein residue contact prediction has attracted more and more attention, particularly with the significant improvement of prediction accuracy in recent years [2, 3]. Theoretically, native residue contacts that are essential for protein structure or function could be inferred from correlated mutations of residue pairs in evolution. With sequence data accumulated at an unprecedented speed, extraction of such coevolution information from multiple sequence alignment (MSA) has become more and more practicable [4-7].

Many early residue contact prediction methods were derived from statistics and information theory, like OMES [8], MI [9], MIp [10] and SCA [11]. However, these methods ignore the transitive correlation between residues and thus generate many false positive results. The inverse covariance matrix and pseudo-likelihood maximization were introduced subsequently to eliminate transitivity in methods such as DCA [12], PSICOV [13], plmDCA [14], GREMLIN [15], CCMpred [16], FreeContact [17] and PconsC2 [18]. These methods effectively reduce false positive predictions by globally considering all inter-residue correlations. More recently, methods like MetaPSICOV [19], SAE-DNN [20], DeepConPred [21], NeBcon [22] and RaptorX-Contact [23] integrated sophisticated machine-learning techniques to further enhance the prediction accuracy. In the latest CASP12 competition, RaptorX-Contact achieved the best performance in the category of template-free modeling targets.

In spite of the general improvement, none of existing methods can attain a robust and steady prediction among all protein targets, mainly because the reliability of coevolution information is guaranteed only when a sufficiently large number of homologous sequences are present in the MSA. Indeed, many protein families lack enough homologous sequences for reliable inference of residue contacts [21], and the predicted residue contact maps of these targets may be dominated by false positives, which hinders the subsequent protein structure prediction/modeling. However, even in the highly noisy residue contact maps for these small-family protein targets, characteristic patterns of specific structural motifs could be identified, because a collective pattern of multiple residue contacts is less likely to be perturbed by individual prediction errors and therefore could be more reliably identified than a single residue contact. Good exemplar structural motifs include parallel and anti-parallel β strands, where consecutive residue pairs from individual β strands establish repetitive contacts in the diagonal and off-diagonal directions on a residue contact map, respectively. Hence, it is possible to identify the residue pairing in interacting β strands from a predicted residue contact map. Identification of β-β pairing would greatly benefit the structural prediction of mainly β proteins, a group of challenging protein targets with complicated topologies. Arguably, structural models of mainly β proteins are reported to be less accurate than those of mainly α proteins, when constructed from residue contact information with comparable levels of accuracies [24].

A great variety of β–β pairing prediction methods have been developed since 1990s [25], including BetaPro [26], MLN/MLN-2S [27], CMM [28] and BCov [29]. Among these methods, the more recent ones, CMM and BCov, make predictions based on coevolution features extracted from the sequence data. Unfortunately, all these previous methods are constructed with the knowledge of native secondary structures and therefore perform unsatisfyingly when fed with predicted secondary structures, which limits their usefulness in practical protein structure prediction. As the first pure predictor modeled without any native structural information, bbcontacts [30] utilizes hidden Markov models to identify β-β pairing from the residue contact map predicted by CCMpred and exhibits a remarkable improvement in performance over all previous algorithms.

Here, we proposed a new approach to predict β-β pairing using ridge detection, a conception that has been well-developed in image processing to capture the axis of an elongated object. Ridge detection was firstly proposed by Haralick [31] in 1983, and was then applied to medical image analysis by Pizer and his co-workers [32, 33]. Lindeberg introduced γ-normalized derivatives and scale-space ridges [34] to better depict the detailed feature of a ridge.

Unlike bbcontacts, in this work, we treated the predicted residue contact map as a raw image and employed the ridge detection to characterize the pattern of consecutive residue contacts for interacting β strands. We designed a multi-stage random forest framework to integrate all ridge-related properties and a number of additional features to predict the β–β contacts. Starting from contact maps predicted by CCMpred [16], our algorithm RDb_2_C (Ridge-Detection-based β-β Contact predictor) shows significant improvements over bbcontacts at both residue and strand levels. Moreover, when connected with the more advanced residue contact predictor RaptorX-Contact [23], RDb_2_C reaches an impressively high level of prediction powers, and the improvement in β–β contact prediction further ameliorates the structure prediction of mainly β proteins.

## Results and Discussion

### Brief introduction of the model

Theoretically, consecutive residue pairs from interacting β strands should present continuous contact points in the diagonal or off-diagonal directions on a native contact map. Even when disguised by prediction noises, the relative strong signals from these β–β contacts are likely to exhibit continuous elongated distributions on a predicted contact map. Here, we adopted the ridge detection, a computer algorithm to identify elongated objects on a 2D image, to capture the characteristic pattern of β-β interactions from predicted contact maps. The ridge information was extracted using the γ-normalized ridge detection method introduced by Lindeberg [34].

Given the original predicted contact map and extracted ridge information, we then developed a novel multi-stage random forest framework to further refine the prediction of β–β contacts. Fig 1 shows the general architecture of the whole algorithm. RDb_2_C starts from a residue contact map predicted based on the amino acid sequence of the target protein, e.g. by CCMpred or by RaptorX-Contact. Besides ridge features, general properties of the input contact map and position of the target residue pair within the map are abstracted as map property features and position features, respectively. The predicted secondary structure probabilities (from DeepCNF [35]) are incorporated as additional features. All features are fed into a 3-stage random forest framework to predict residue pairing in interacting β strands.

**Fig 1.**
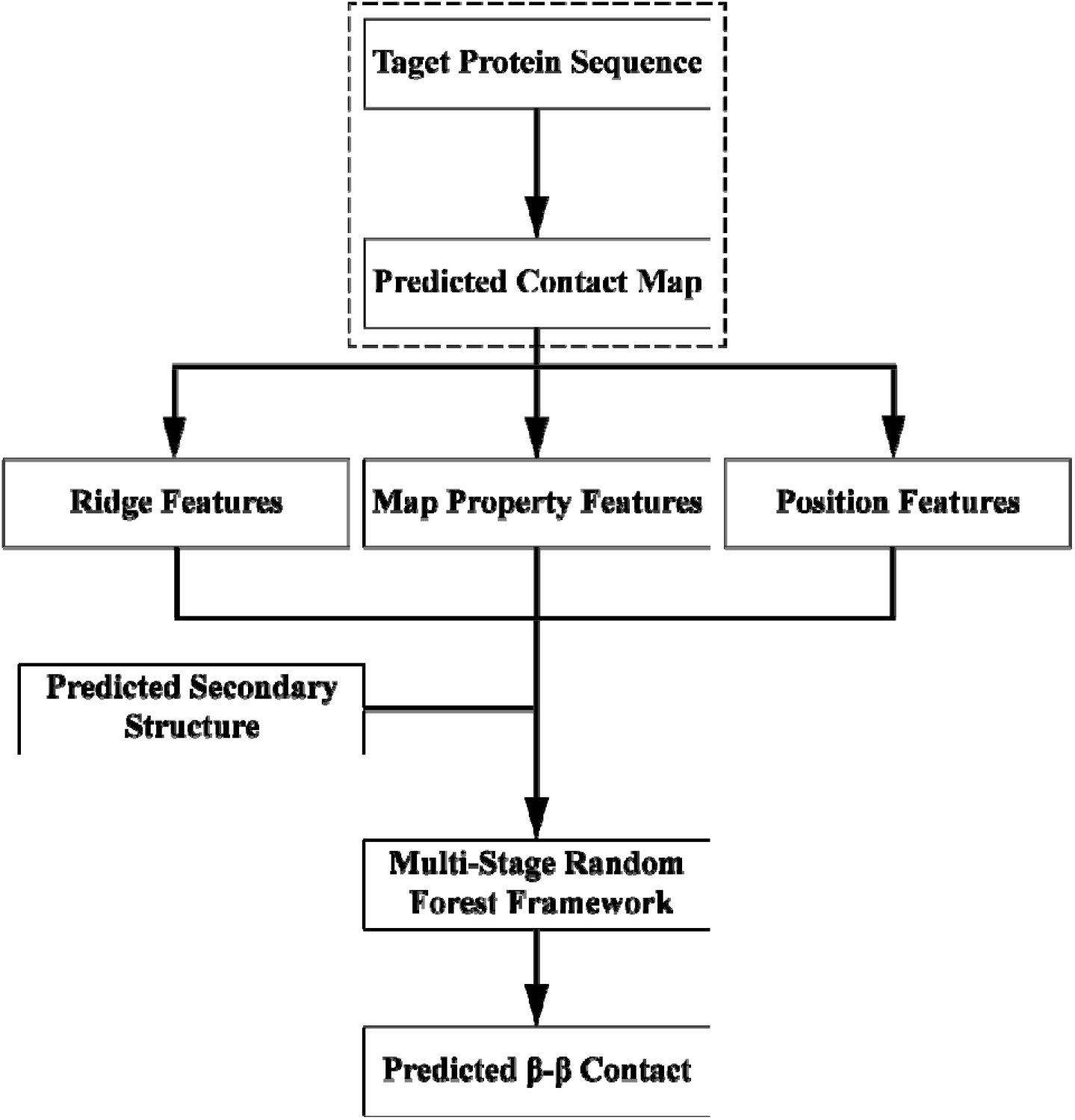
The general flow chart of RDb_2_C.

Specifically, we constructed 4 random forest models with different window sizes (3×3, 5×5, 7×7 and 9×9) at the first stage. The prediction results of the first stage models were then combined in the second stage and further optimized in the third stage by taking the preceding-stage results as input features. The model optimization of each stage was performed using 5-fold cross-validation on a training set containing 493 proteins. Further testing and performance evaluation were conducted on two well-established datasets, BetaSheet916 [26] and BetaSheet1452 [29]. Notably, redundancy between the training and test datasets has been carefully removed.

### Performance evaluation of the model

The performance of RDb_2_C models at all stages was evaluated in the cross-validation as well as the BetaSheet916 and BetaSheet1452 test sets. Table 1 summarizes the residue-level performance. Clearly, all models show robust and balanced performance between the two independent test sets, which indicates appropriate model training. It is noticeable that cross-validation exhibits lower F1-scores than the test sets. This difference may be attributed to the presence of more small-family proteins in the training set than in the test sets (Fig 2): 18.05% of the training set proteins have less than *L* sequences in the MSA (*L* is the protein length), whereas the percentage reduces to only 7.21% and 1.31% in the BetaSheet916 and BetaSheet1452 sets, respectively.

**Table 1.**
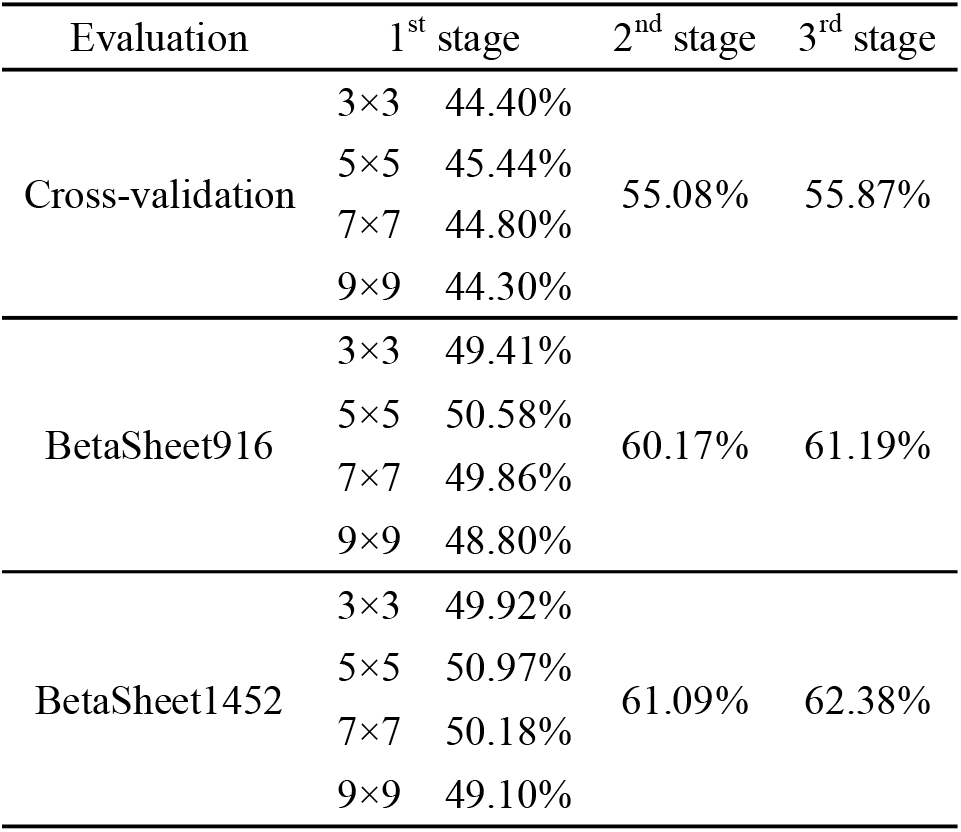
Residue-level F1-scores of all models in the 5-fold cross-validation, BetaSheet916 and BetaSheet1452 sets.

**Fig 2.**
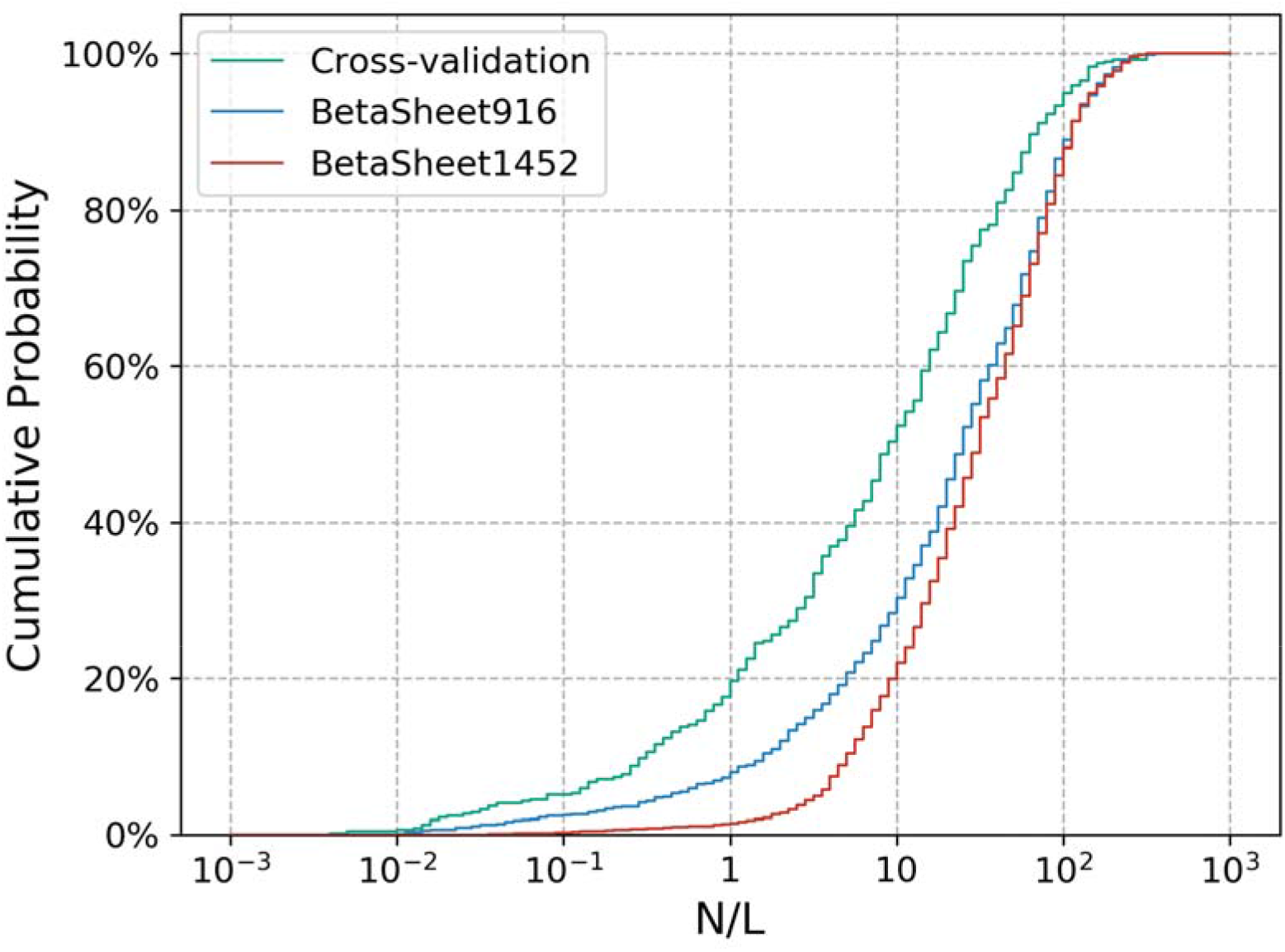
The cumulative distributions for training and test sets with the respect of *N*/*L*. *N* is the number of sequences in the MSA and *L* is the protein length. There are more proteins in the training set with limited numbers of homologous sequences (*N*/*L* < 1) than in the BetaSheet916 and BetaSheet1452 sets.

The first-stage models attain the optimal performance at the window size of 5 in both cross-validation and test sets. We suspect that the larger windows include more useful information but also introduce more noises that eventually impairs the model performance, and that balance of useful information and noise may be achieved at the window size of 5. However, models constructed at various window sizes could provide complementary information. Accordingly, the second-stage models that combine information achieved at all window sizes exhibit significant improvement (~10 percentage points) in F1-scores over the first-stage ones. At the third stage, further optimization slightly improves the F1-score to 61.19% and 62.38% in the BetaSheet916 and BetaSheet1452 sets, respectively.

To justify the effectiveness of novel features we proposed in this work, we evaluated the feature importance for all first-stage models. The feature importance was evaluated by re-conducting the model optimization and cross-validation without the corresponding features. As shown in Table 2, all features are essential for the model, since removal of each type weakens the performance. Moreover, all first-stage models exhibit a uniform trend: the ridge features and the original CCMpred map jointly make the major contribution to the prediction power (see the loss of >20 percentage points after removal of both features). Although the ridge features are derived from the CCMpred map, removing ridge features alone significantly deteriorates the F1-score, especially for models of small window sizes, possibly because these features are capable of summarizing the local information and depicting the local shape character of a predicted contact map. Therefore, the ridge features introduced in this work effectively capture the residue contact pattern of β-β interactions. In addition, the secondary structure information predicted by DeepCNF is also constructive to our model, which is reasonable considering that proper assignment of β residues are the prerequisite for the prediction of β-β contacts.

**Table 2.**
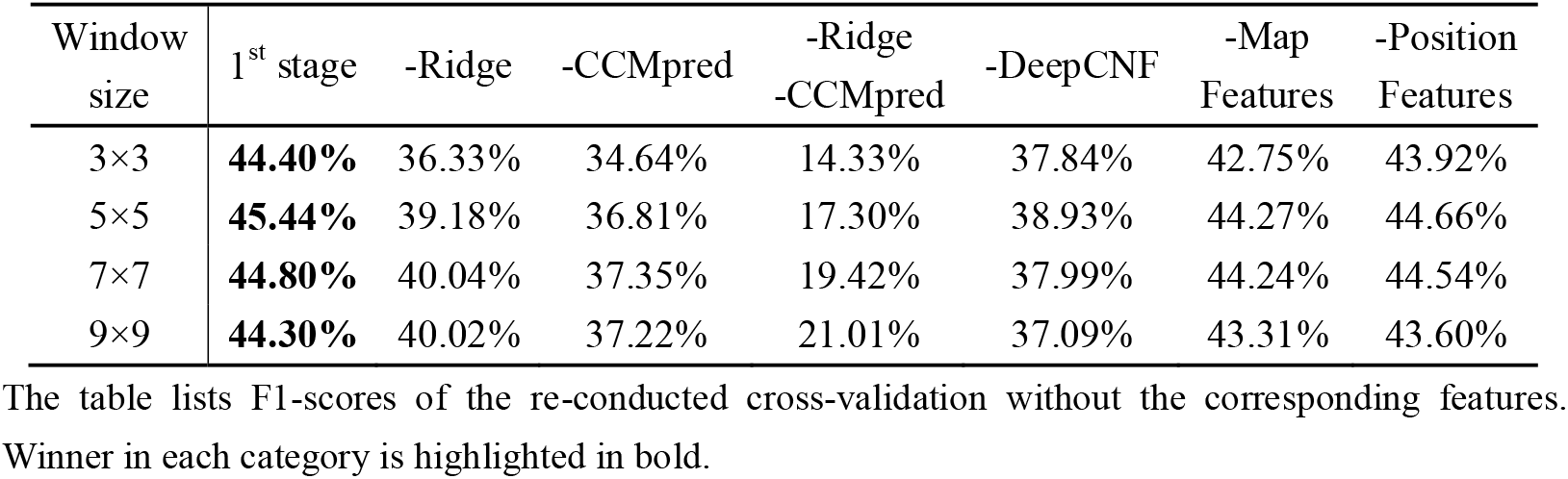
The feature importance in the first-stage models.

As expected, when using the native secondary structures assigned by DSSP [36] instead of the predicted ones as input, the DSSP-based models provide improvement of ~10 percentage points to the residue-level predictions (Table 3). Thus, more accurate secondary structure prediction algorithm could further improve the performance potentially. Table 4 summarizes the strand-level performance in the BetaSheet916 and BetaSheet1452 sets. Notably, the strand-level performance was only evaluated using the DSSP-based framework due to the requirement of exact secondary structure information in the assignment of β strands. Similar to residue-level results (see Table 1), the strand-level models are progressively refined with stages, with the final F1-scores reaching 75.40% and 76.55% in the BetaSheet916 and BetaSheet1452 sets, respectively.

**Table 3.** Residue-level performance of RDb2C constructed with DeepCNF-predicted and DSSP-assigned secondary structure information.

| Secondary structure | Models | BetaSheet916 |  |  | BetaSheet1452 |  |  |
| --- | --- | --- | --- | --- | --- | --- | --- |
|  |  | Precision | Recall | F1-score | Precision | Recall | F1-score |
| Predicted | 1<sup>st</sup> stage | 63.94% | 41.84% | 50.58% | 57.61% | 45.71% | 50.97% |
|  | 2<sup>nd</sup> stage | 65.03% | **55.99%** | 60.17% | 64.50% | **58.02%** | 61.09% |
|  | 3<sup>rd</sup> stage | **68.00%** | 55.62% | **61.19%** | **67.91%** | 57.69% | **62.38%** |
| DSSP | 1<sup>st</sup> stage | 69.92% | 49.94% | 58.26% | 62.71% | 54.22% | 58.16% |
|  | 2<sup>nd</sup> stage | 75.79% | 64.00% | 69.40% | 75.74% | 66.07% | 70.58% |
|  | 3<sup>rd</sup> stage | **76.28%** | **65.94%** | **70.74%** | **76.56%** | **67.86%** | **71.95%** |

Performances of the models with the window size of 5 are listed here as the representatives of the first-stage models. Winner in each category is highlighted in bold.

**Table 4.**
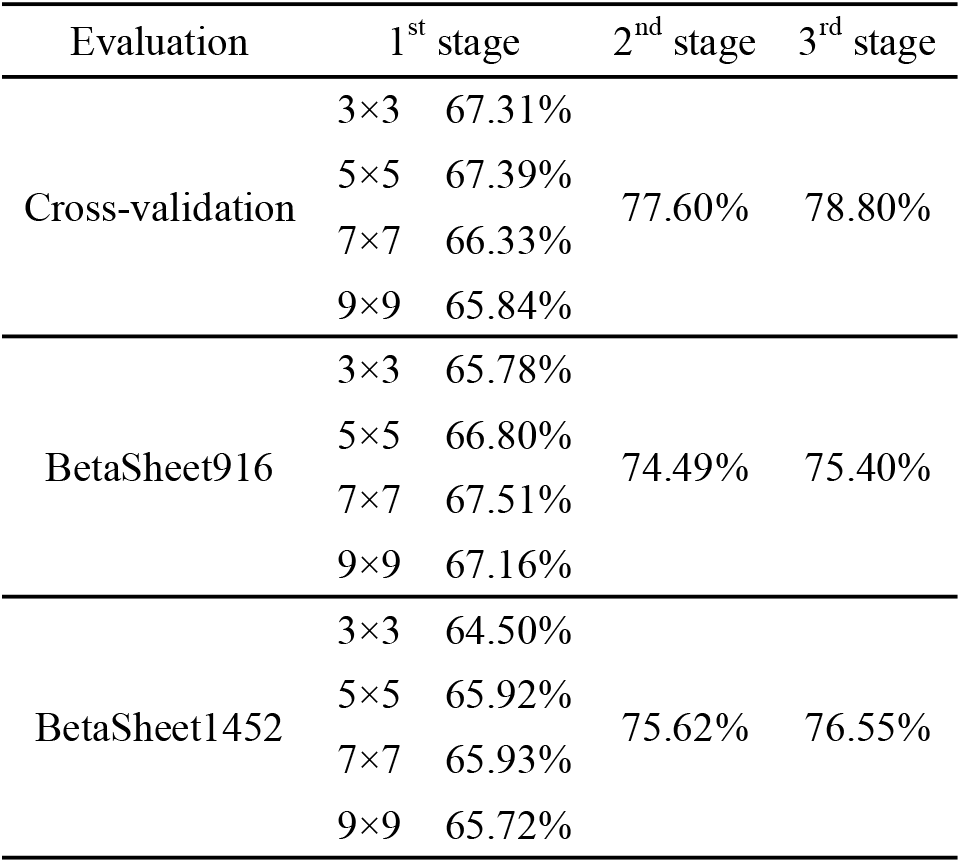
Strand-level F1-scores of all models in the 5-fold cross-validation, BetaSheet916 and BetaSheet1452 sets.

### Comparison with bbcontacts

Here, we mainly compared RDb_2_C with bbcontacts, the best predictor so far among all previous methods. The performance of RDb_2_C and bbcontacts could be fairly compared since both methods take CCMpred contact maps as input. Fig 3 presents the Precision-Recall (PR) curves of RDb_2_C and bbcontacts at the residue and strand levels in the BetaSheet916 and BetaSheet1452 sets, respectively. At the residue level, RDb_2_C outperforms bbcontacts on the whole range, especially in the region of high-Precision values. Specifically, with the sacrifice of Recall, RDb_2_C could approach the Precision level of 90-100%, which means that top-scored predictions of RDb_2_C are almost error-less and thus can be directly applied to practical structure prediction. In contrast, bbcontacts can only access the Precision level of 70-80%. As for the strand-level results, despite the crossing of PR curves, RDb_2_C outperforms bbcontacts in most ranges, particularly at the high-Precision region that reflects the quality of top-scored predictions.

**Fig 3.**
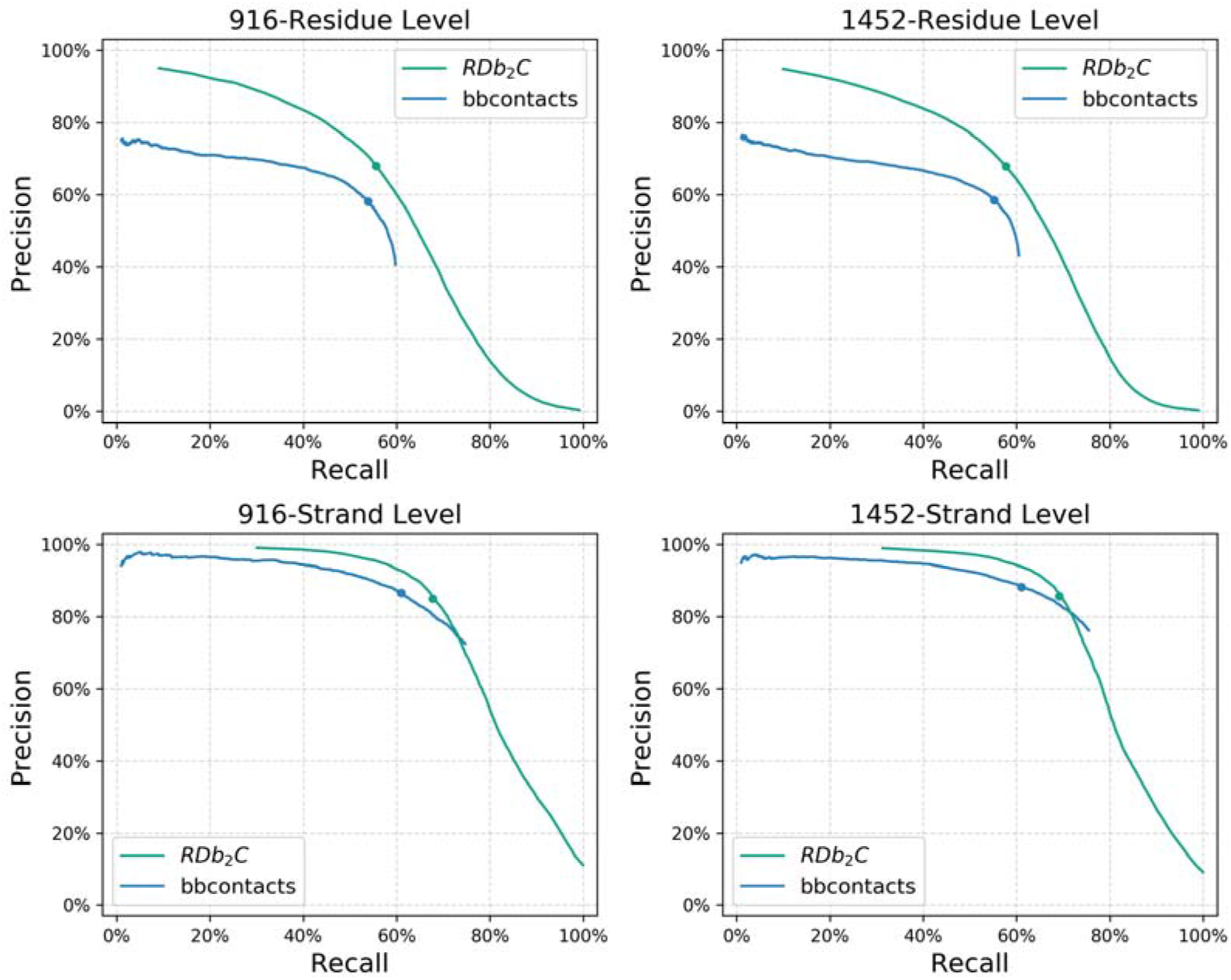
The PR curves in the BetaSheet916 and BetaSheet1452 sets. The comparison is shown for RDb_2_C (green) and bbcontacts (blue), at the residue level (top row) and strand level (bottom row) as well as in the BetaSheet916 (left column) and BetaSheet1452 (right column) sets, respectively. Performances at the suggested cutoffs are marked as dots on the PR curves.

Detailed comparison of the two methods at their respective suggested cutoffs is listed in Table 5. Both RDb_2_C and bbcontacts are quite robust between the BetaSheet916 and BetaSheet1452 sets. In comparison to the reported numbers in the original paper, performance of bbcontacts increases substantially (residue-level F1-score of ~56% *vs.* ~50% in the paper), possibly due to the enhanced prediction accuracy of CCMpred with the accumulation of sequence data in the past years. However, RDb_2_C still outperforms bbcontacts by ~6 percentage points at the residue level, in terms of F1-scores. At the strand level, RDb_2_C and bbcontacts have different preferences of Precision and Recall, but comprehensively RDb_2_C achieves a higher level of F1-scores (~76%) and outperforms bbcontacts by ~4 percentage points.

**Table 5.**
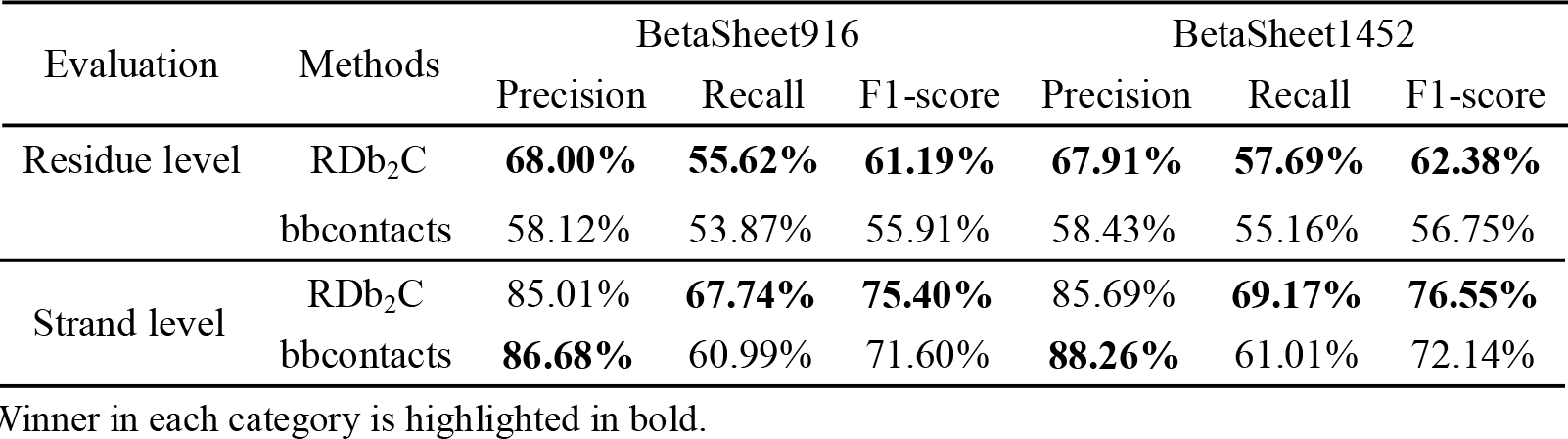
Performance comparison of RDb_2_C and bbcontacts at residue level and strand level.

Subsequently, we systematically compared the F1-scores of RDb_2_C and bbcontacts for individual proteins in the BetaSheet916 and BetaSheet1452 sets (Fig 4). At the residue level, RDb_2_C outperforms bbcontacts on 69.32% targets of the BetaSheet916 set and 72.56% targets of the BetaSheet1452 set, respectively, in terms of F1-scores. The superiority of RDb_2_C over bbcontacts is statistically significant (p-value < 10^-10^) in both test sets. At the strand level, RDb_2_C exhibits better performance on 61.57% and 63.36% targets of the BetaSheet916 and BetaSheet1452 sets, respectively, and this advantage is also statistically significant with p-values < 10^-10^.

**Fig 4.**
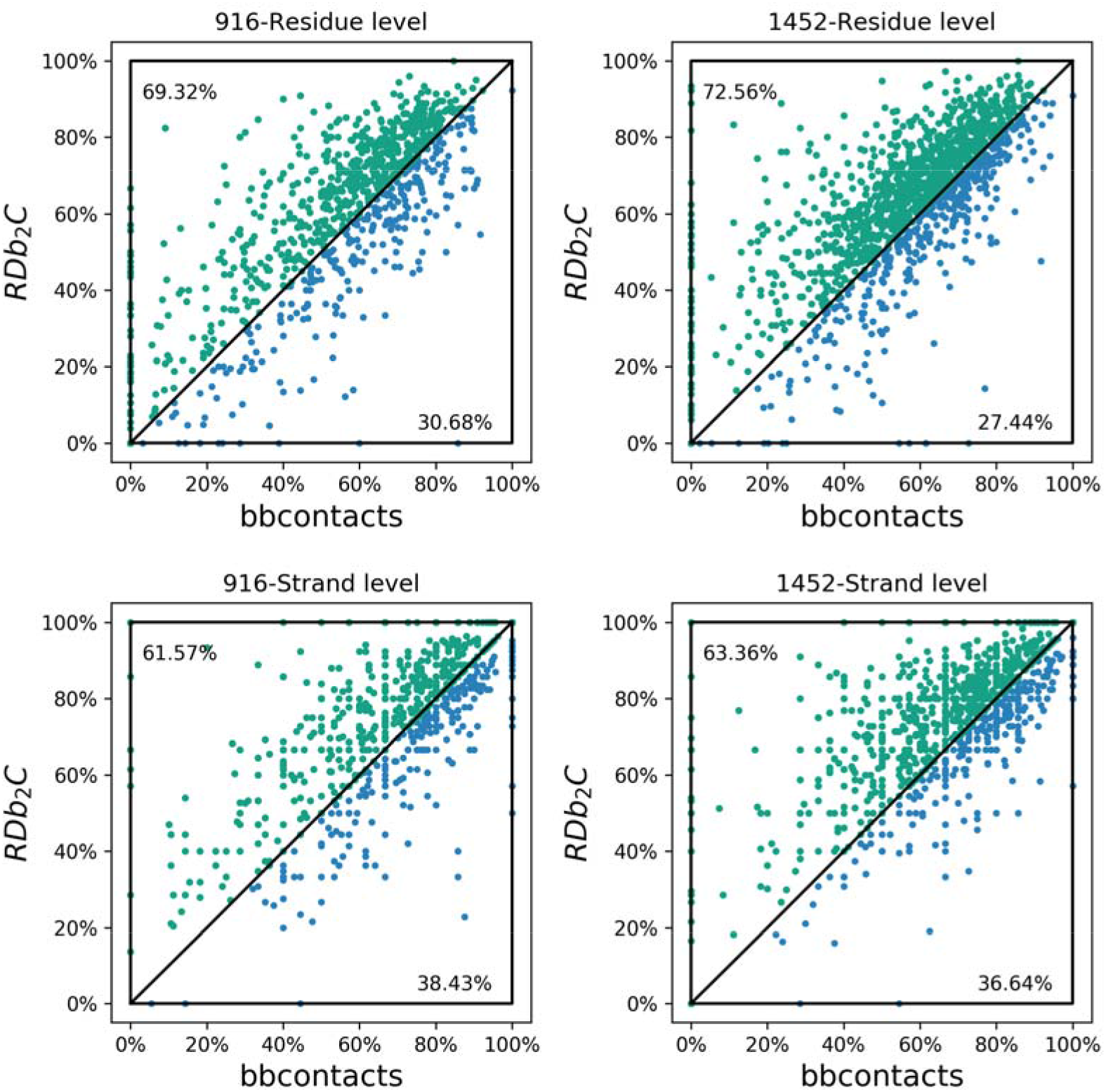
Comparison of RDb_2_C and bbcontacts for individual proteins of the BetaSheet916 and BetaSheet1452 sets. Each individual protein is represented as a dot. The green dots and blue dots represent targets that are better predicted by RDb_2_C and by bbcontacts, respectively, in terms of F1-scores. Tie cases are bisected to two methods. In both test sets and at both residue and strand levels, RDb_2_C outperforms bbcontacts significantly (p-value < 10^-10^).

To compare with other previous methods that have reported results only for DSSP-based predictions, we evaluated the DSSP-based models for RDb_2_C and bbcontacts at the residue level. As shown in Table 6, RDb_2_C outperforms bbcontacts by 2-3 percentage points with the knowledge of native secondary structures, while both RDb_2_C and bbcontacts remarkably outperform previous methods by large margins.

**Table 6.**
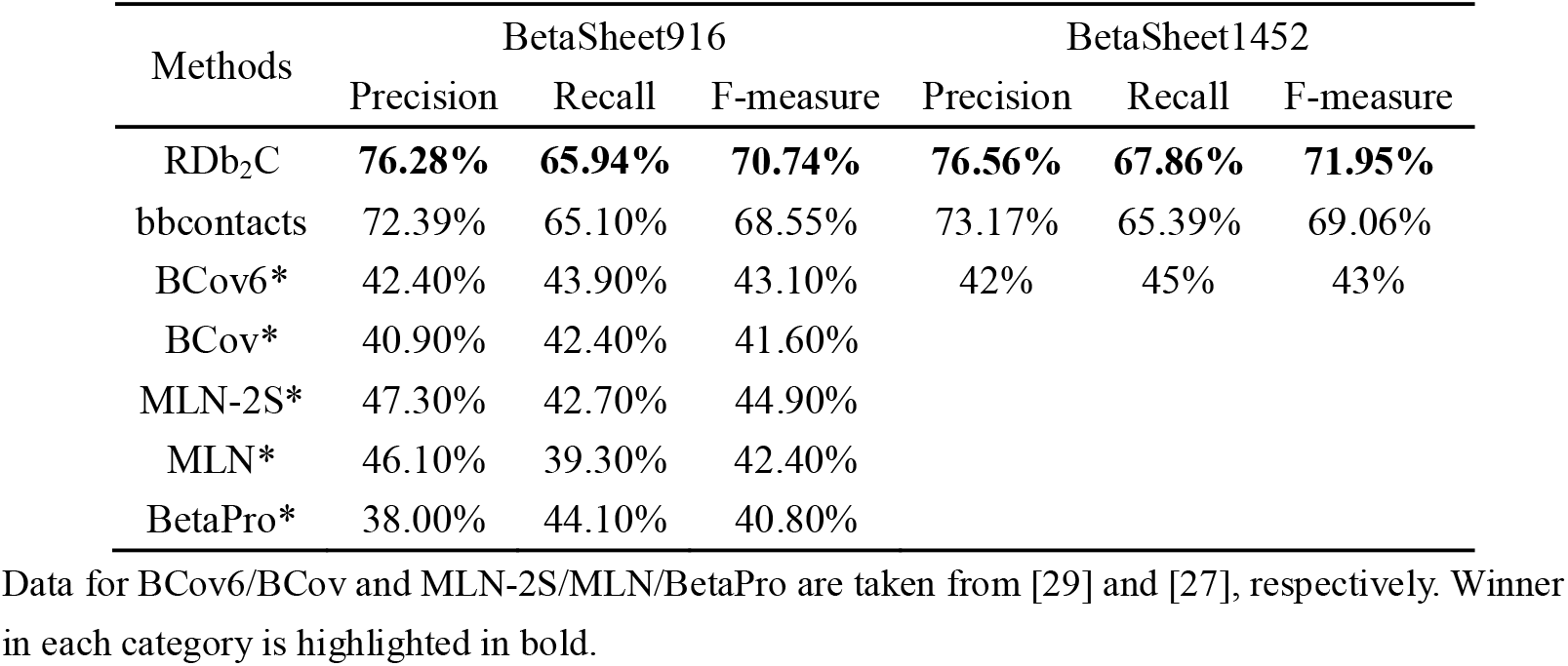
Performance comparison of DSSP-based RDb_2_C, bbcontacts and other methods at the residue level.

The advantage of RDb_2_C over bbcontacts in models constructed with predicted secondary structures may arise from two facets of differences: 1) different programs adopted for secondary structure prediction (DeepCNF in RDb_2_C *vs.* PSIPRED pipelined with HHsuite in bbcontacts); 2) difference in program design. To test the former point, we first compared the prediction power of DeepCNF and the PSIPRED pipeline used in bbcontacts (Table 7). In all categories, DeepCNF has comparable or slightly weaker prediction power than the PSIPRED pipeline. Furthermore, we tested the bbcontacts model constructed with DeepCNF prediction as input. The DeepCNF-based bbcontacts model achieves residue-level F1-scores of 55.17% and 56.19% in the BetaSheet916 and BetaSheet1452 sets, respectively, nearly indistinguishable with the original PSIPRED-based model (55.91% and 56.75%, respectively). Therefore, the superiority of RDb_2_C over bbcontacts is mainly attributed to the unique design of our method, for instance, the application of ridge detection and the novel multi-stage framework.

**Table 7.**
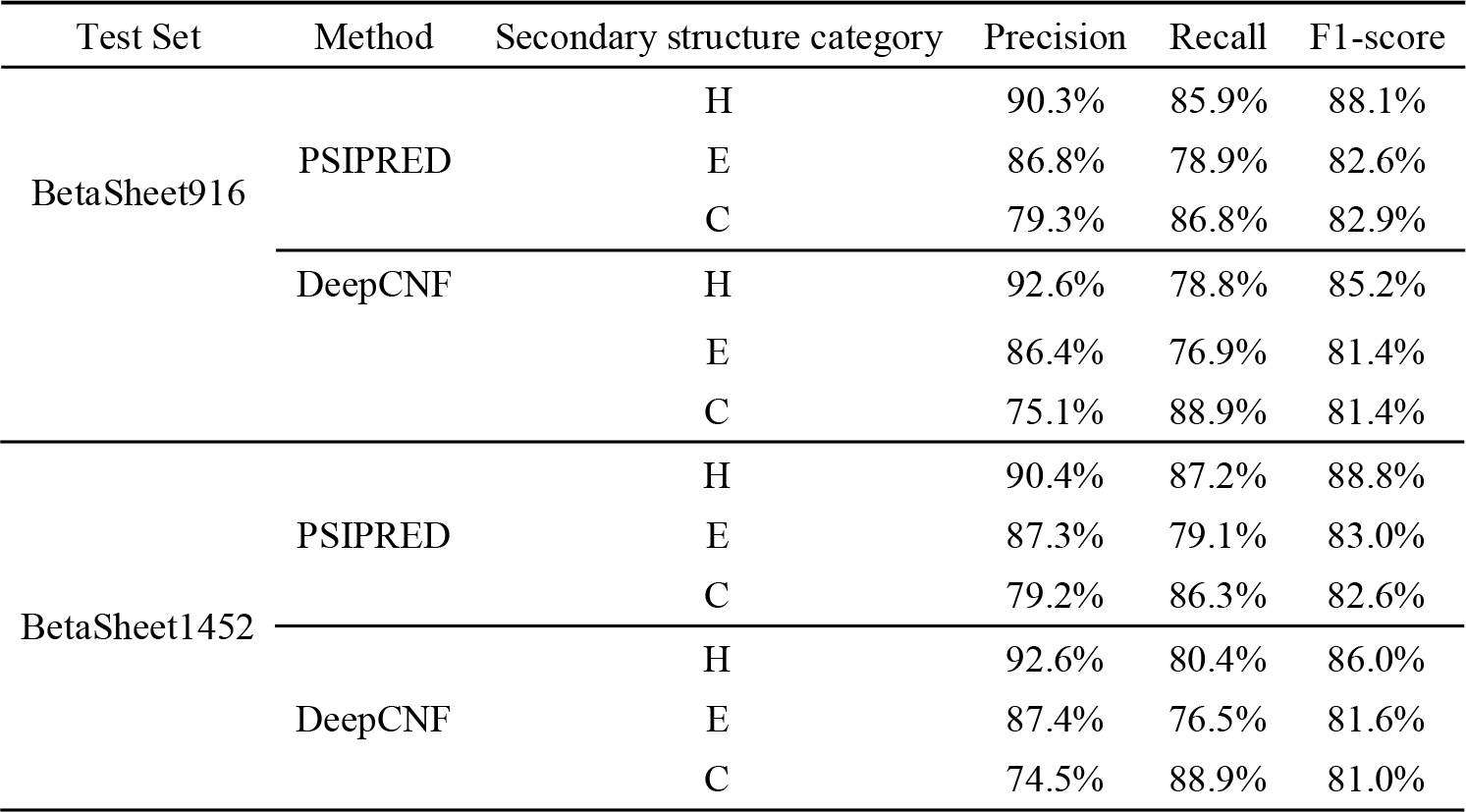
Performance comparison of DeepCNF and PSIPRED in the BetaSheet916 and BetaSheet1452 sets.

In Fig 5, we include three protein cases as examples to show the improvement in the prediction of β-β contacts using RDb_2_C and bbcontacts. In these examples, the raw CCMpred maps are dominated by noises, which hinders visual identification of β-β interactions. Although both RDb_2_C and bbcontacts are capable of finding signals from the noises, the native β-β contacts could be more successfully identified by RDb_2_C, at both residue and strand levels.

**Fig 5.**
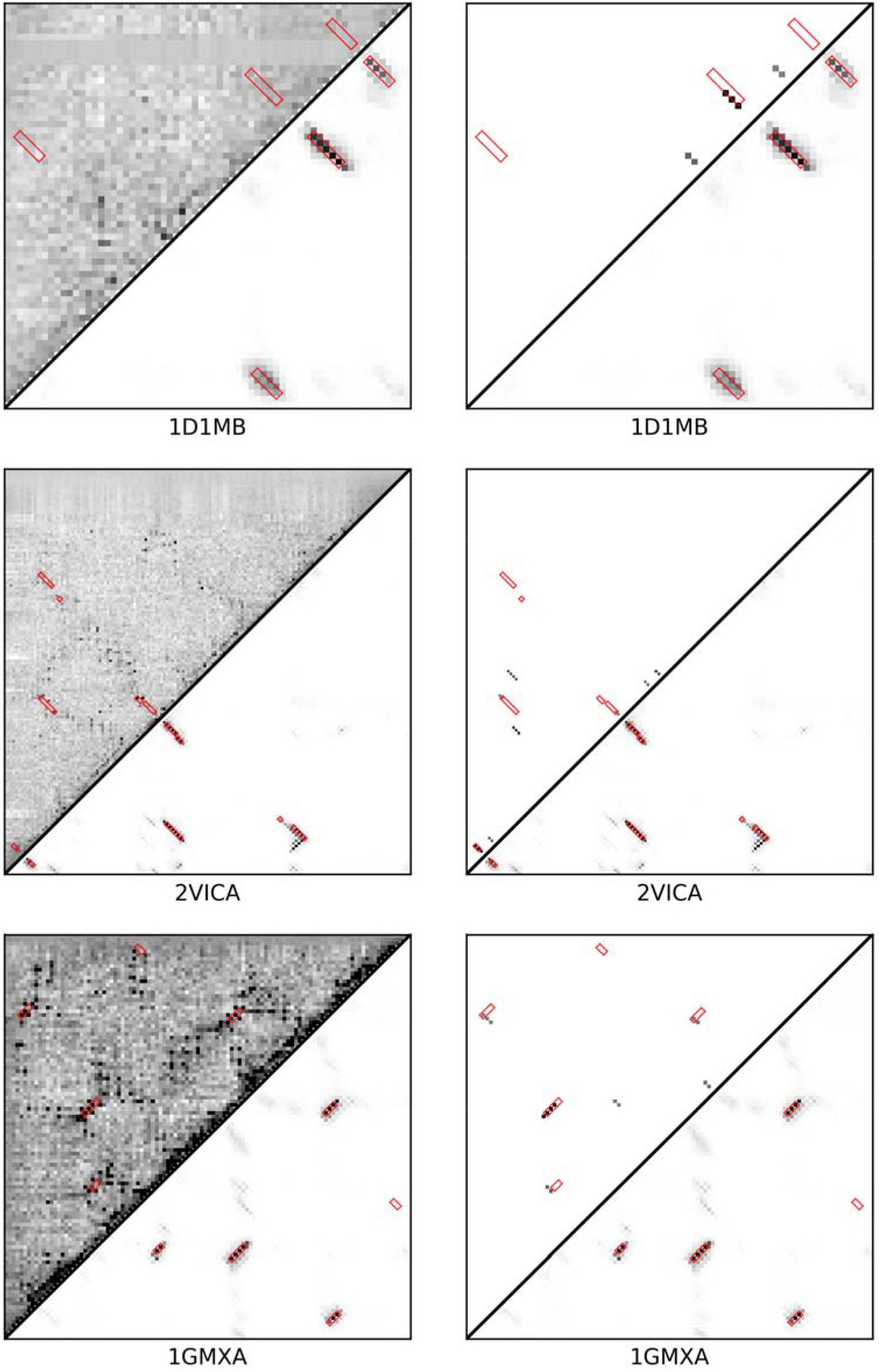
Case studies for CCMpred-based predictions. We illustrate three CCMpred-based case studies. In the left-handed panel, the upper left triangle is the raw CCMpred map, while the lower right triangle is the prediction by RDb_2_C. In the right-handed panel, the upper left triangle is replaced by results of bbcontacts to facilitate direct comparison with RDb_2_C (i.e. the lower right triangle). The native β-β contact regions are highlighted by red boxes.

### Pipelined with RaptorX-Contact

RDb_2_C is developed to refine the prediction of β-β contacts from any predicted contact maps. To verify the general applicability, we tested the performance of our method on contact maps predicted by RaptorX-Contact, one of the most successful residue contact predictors in the latest CASP12 competition. The whole framework was optimized in the same training set, except that the raw maps were obtained from the RaptorX-Contact server. Due to the failure in processing a few protein targets by the server, available proteins in the training set reduces to 383 CATH domains (S1 Table). Considering the time consumption in server submission, this test was conducted only on the BetaSheet916 set. Similarly, the number of available proteins in the BetaSheet916 set was shrunk to 858.

To evaluate the prediction powers of RaptorX-Contact and CCMpred in the β regions, we collected the prediction scores of all pairs of β residues as referred by DSSP assignment. These scores were then sorted and an adjustable cutoff value was used to identify the positive predictions. In this manner, Precision and Recall values at various cutoff values could be collected, which enables the plotting of PR curve as well as the calculation of optimal F1-score. Noticeably, the F1-scores derived in this way may be overestimated, because knowledge of native secondary structures is utilized and because the cutoff is self-optimized rather than estimated independently. Results suggest that RaptorX-Contact provides significantly more accurate residue contact prediction than CCMpred. As for β-β contacts, CCMpred only achieves an F1-score of 20.28%, while RaptorX-Contact attains 60.23%. However, even starting from the poor contact maps of CCMpred, RDb_2_C could improve the prediction of β-β contacts to a level comparable to RaptorX-Contact (~61%, see Table 3).

The evaluation of our models optimized on the RaptorX-Contact maps is summarized in Table 8. Unlike previous results (see Table 1), the model performance shows negligible improvement in sequential stages, which indicates that prediction could terminate in early stages when the input residue contact maps are of high quality. Nevertheless, RDb_2_C finally reaches impressively high F1-scores of 76.17% and 85.65% at the residue and strand levels, respectively. Notably, performance of these levels could ensure both prediction accuracy (Precision) and coverage of native β-β contacts (Recall) at sufficiently high values (>70%), which thus would greatly benefit the tertiary structure prediction of mainly β proteins.

**Table 8.**
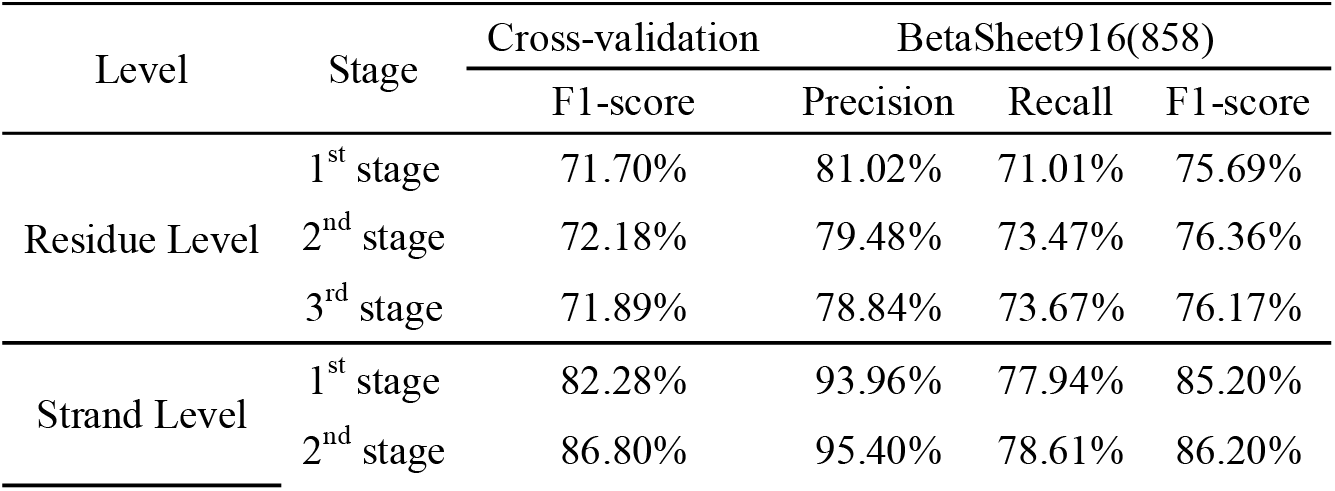
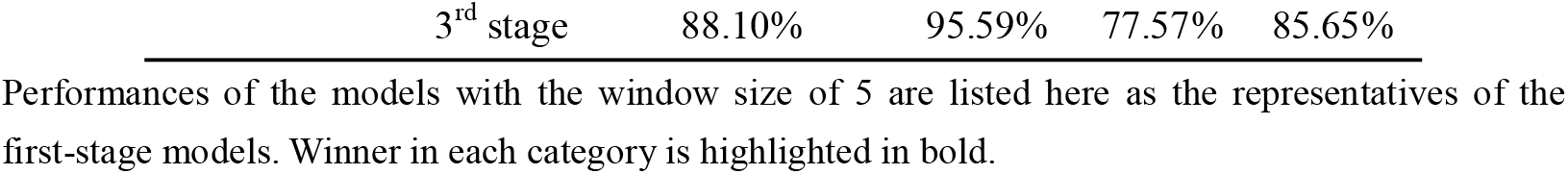
Performance of RDb_2_C at residue level and strand level on the 5-fold cross-validation and shrunk BetaSheet916 set.

| Level | Stage | Cross-validation | BetaSheet916(858) |  |  |
| --- | --- | --- | --- | --- | --- |
|  |  | F1-score | Precision | Recall | F1-score |
| Residue Level | 1<sup>st</sup> stage | 71.70% | 81.02% | 71.01% | 75.69% |
|  | 2<sup>nd</sup> stage | 72.18% | 79.48% | 73.47% | 76.36% |
|  | 3<sup>rd</sup> stage | 71.89% | 78.84% | 73.67% | 76.17% |
| Strand Level | 1<sup>st</sup> stage | 82.28% | 93.96% | 77.94% | 85.20% |
|  | 2<sup>nd</sup> stage | 86.80% | 95.40% | 78.61% | 86.20% |

|  |  |  |  |  |
| --- | --- | --- | --- | --- |
| 3<sup>rd</sup> stage | 88.10% | 95.59% | 77.57% | 85.65% |

Performances of the models with the window size of 5 are listed here as the representatives of the first-stage models. Winner in each category is highlighted in bold.

In comparison to CCMpred-based results (see Table 5), F1-scores are improved by ~15 percentage points, which is mainly attributed to the greatly enhanced quality of residue contact map predicted by RaptorX-Contact. As suggested by the evaluation of feature importance (Table 9), ridge features and raw RaptorX-Contact scores in combination still provide major contribution to the prediction power. However, with the remarkable improvement in the quality of the input map, contribution of the individual ridge features becomes less important, when compared with CCMpred-based predictions (see Table 2).

**Table 9.**
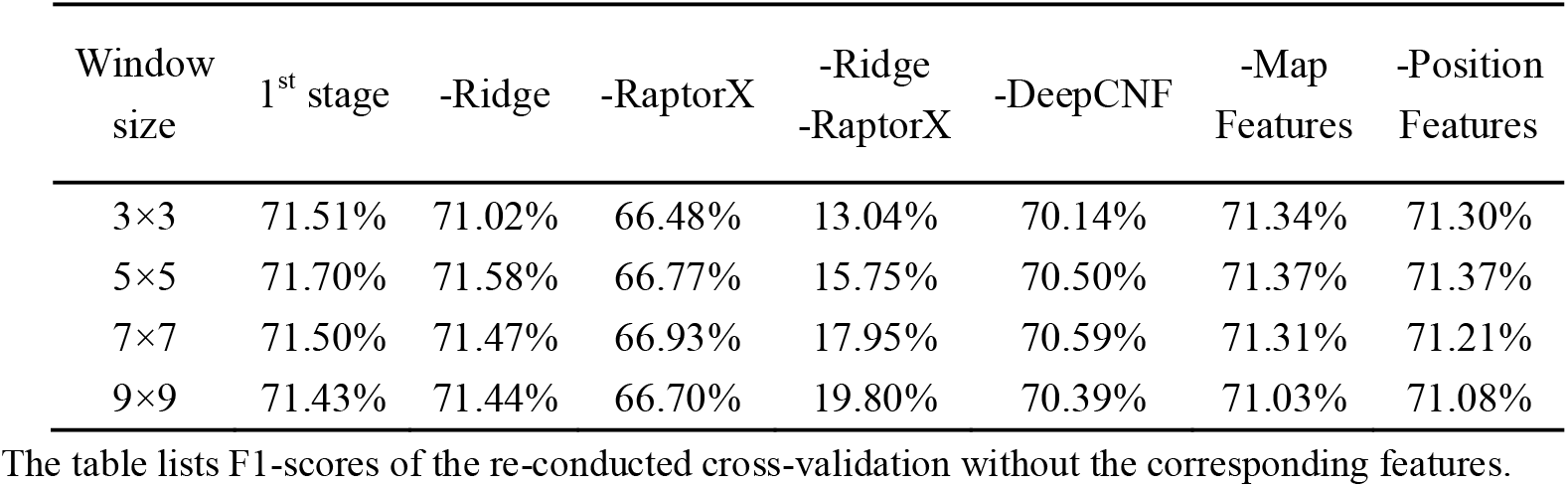
The feature importance in the first-stage models starting with RaptorX-Contact predictions.

On the other hand, RDb_2_C is capable of further improving the high-quality contact prediction of RaptorX-Contact. In specific, the F1-score of β-β contacts increases from an estimated number of ~60% to 76.17%. The great improvement by RDb_2_C is also illustrated in the PR curves (Fig 6). Considering that knowledge of native secondary structures is required in the generation of RaptorX-Contact curve, we also included the PR curve of the DSSP-based RDb_2_C model for a fair comparison. The DSSP-based RDb_2_C model could further improve F1-score to 85.30%. Fig 7 shows the comparison of RDb_2_C over RaptorX-Contact on two protein cases, where the raw RaptorX-Contact maps are noisy but native β-β contacts could be successfully recognized after refinement using RDb_2_C.

**Fig 6.**
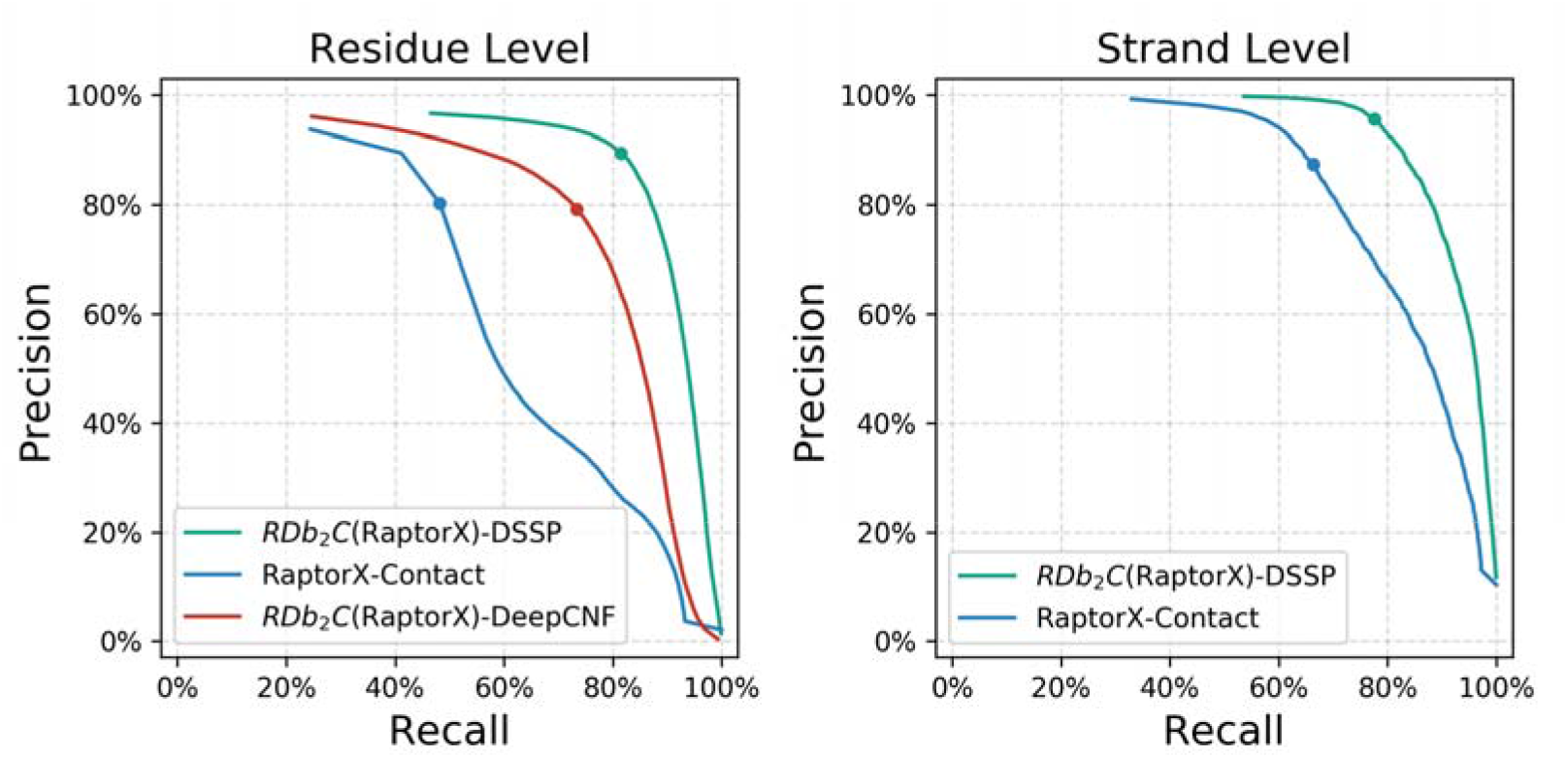
The PR curves in the shrunk BetaSheet916 set. RDb_2_C (green for DSSP-based model and red for DeepCNF-based model) exhibits significant improvement over the raw RaptorX-Contact prediction (blue). The dots on the PR curve illustrate model performance at the suggested RDb_2_C cutoffs and the optimized RaptorX-Contact cutoffs.

**Fig 7.**
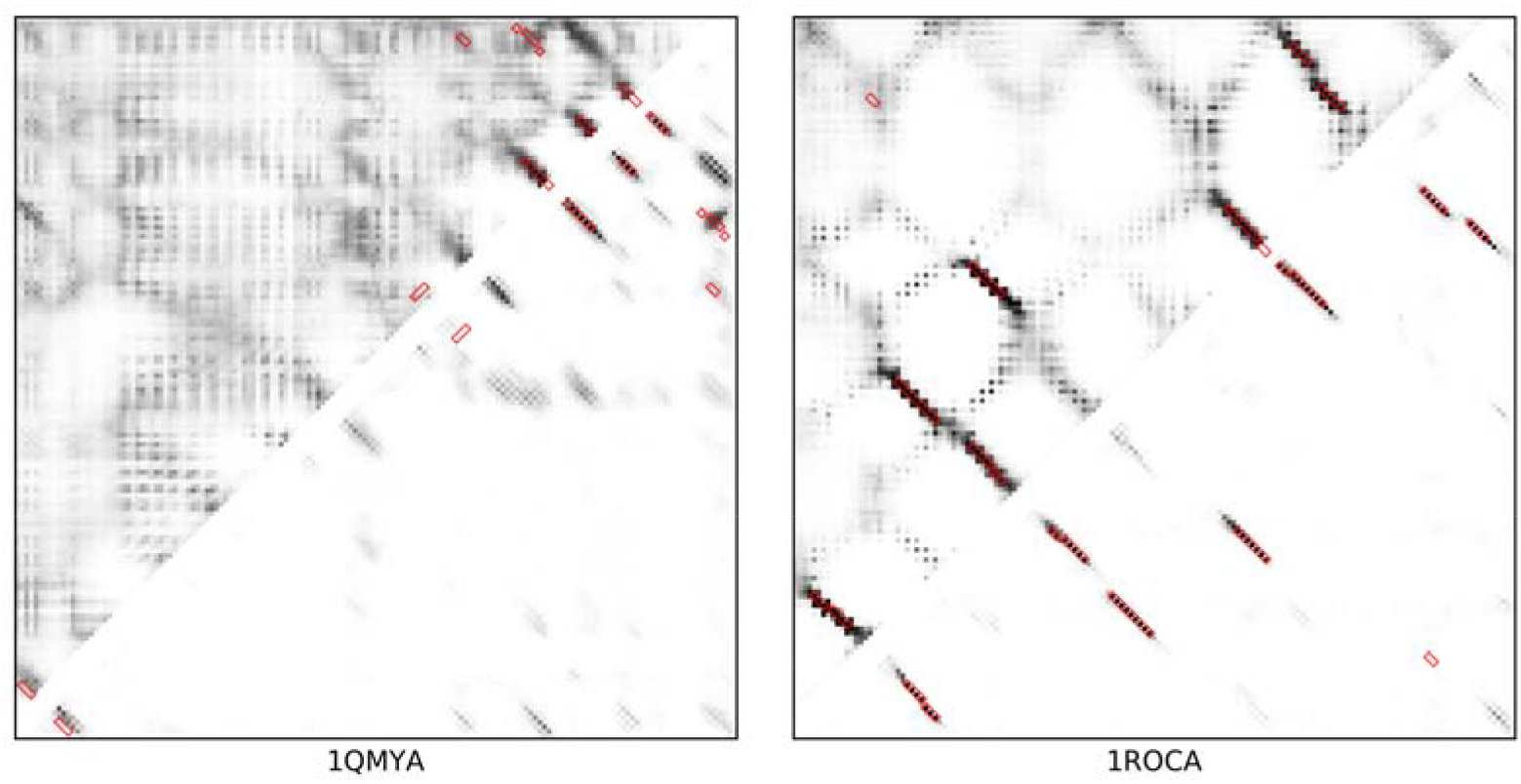
Case studies for RaptorX-Contact-based predictions. We illustrate two RaptorX-Contact-based case studies: 1QMYA (left) and 1ROCA (right). In each plot, the upper left triangle is the raw RaptorX-Contact map, while the lower right triangle is the prediction by RDb_2_C. The native β-β contact regions are highlighted by red boxes.

### Evaluation for the contribution in tertiary structure prediction

In order to justify the effectiveness of our method in the practical structure prediction, we chose 61 mainly β proteins (with ≥50% of β residues) from the shrunk BetaSheet916 set (S2 Table) and constructed the tertiary structure models of them with predicted contacts taken as constraints, following the standard CONFOLD protocol [37]. As the numbers of predicted and native β-β contact pairs are always less than 0.5*L* (S2 Table; *L* is the protein length), which is not sufficient for structural modeling, we retained all β-β contacts predicted by the RDb_2_C model in pipeline with RaptorX-Contact at the suggested cutoff as the highly reliable contact pairs, and then enriched the list of contact pairs to 1*L* by collecting the high-ranked and non-redundant RaptorX-Contact predictions. These top 1*L* residue contacts were used as distance restraints to fold the protein. Specifically, a strict restraint of 3.5-6Å was applied to constrain the C_β_ atoms of residue pairs from the more reliable RDb_2_C prediction, whereas a loose restraint of 3.5-10Å was adopted for the non-redundant residue pairs enriched from RaptorX-Contact results because of their lower confidence level. As a control, the top 1*L* residue contacts were directly chosen from the RaptorX-Contact prediction and a uniform standard restraint of 3.5-8Å was engaged to constrain the C_β_ atoms of these residue pairs.

For each tested protein, the model with the best TM-score [38] within the top 5 models reported by CONFOLD was chosen for evaluation. According to our results, models constructed with the top 1*L* RaptorX-Contact predictions reach an average TM-score of 0.442. In contrast, when supplemented with the refined top 1*L* contacts by RDb_2_C, the average TM-score markedly increases to 0.506. Specifically, among the 61 mainly β proteins, prediction using RDb_2_C refinement outperforms that using RaptorX-Contact raw scores in 83.61% and 85.25% of cases when evaluated by TM-score and RMSD, respectively (Fig 8 and S2 Table). The superiority of RDb_2_C over RaptorX-Contact is statistically significant (p-value < 10^-8^) for both RMSD and TM-score.

**Fig 8.**
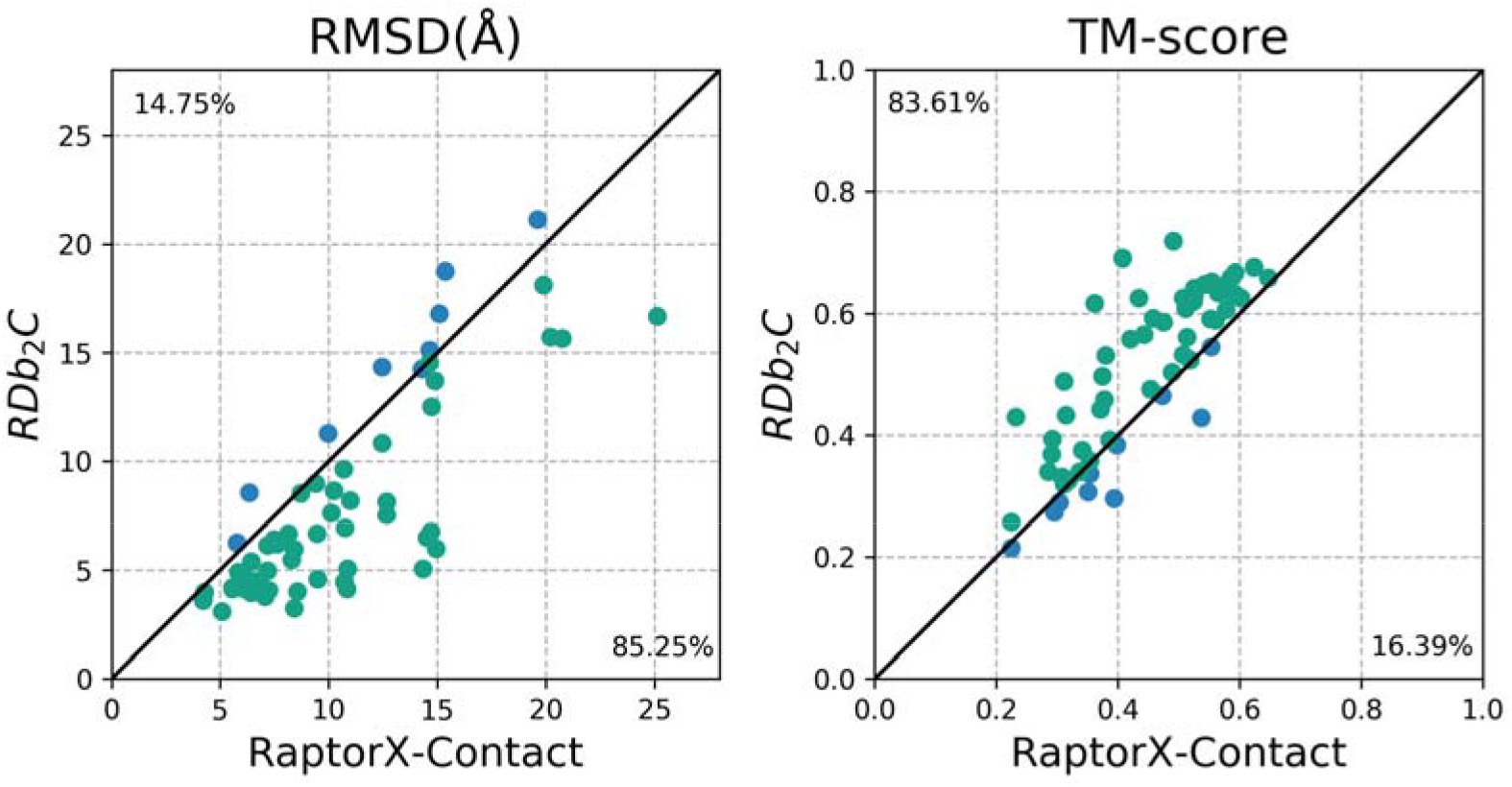
Comparison of the best of the top 5 models generated using the RaptorX-Contact prediction and the RDb_2_C refinement for individual targets of the 61 mainly β proteins. The green dots and blue dots represent targets that are better predicted by RDb_2_C and by RaptorX-Contact respectively. Detailed results are listed in S2 Table. For both RMSD and TM-score, RDb_2_C outperforms RaptorX-Contact significantly (p-value < 10^-8^).

Fig 9 shows the comparison of one protein case, where the RDb_2_C results successfully correct the topology mismatch in the RaptorX-Contact model. Because our predictions focus on the more detailed hydrogen bonding interactions, instead of direct use as the distance restraints for residue C_β_ atoms, it is possible to further improve the structure prediction by utilizing our prediction more delicately, for instance, to restrain the respective hydrogen bonding donors and acceptors of two paired β residues.

**Fig 9.**
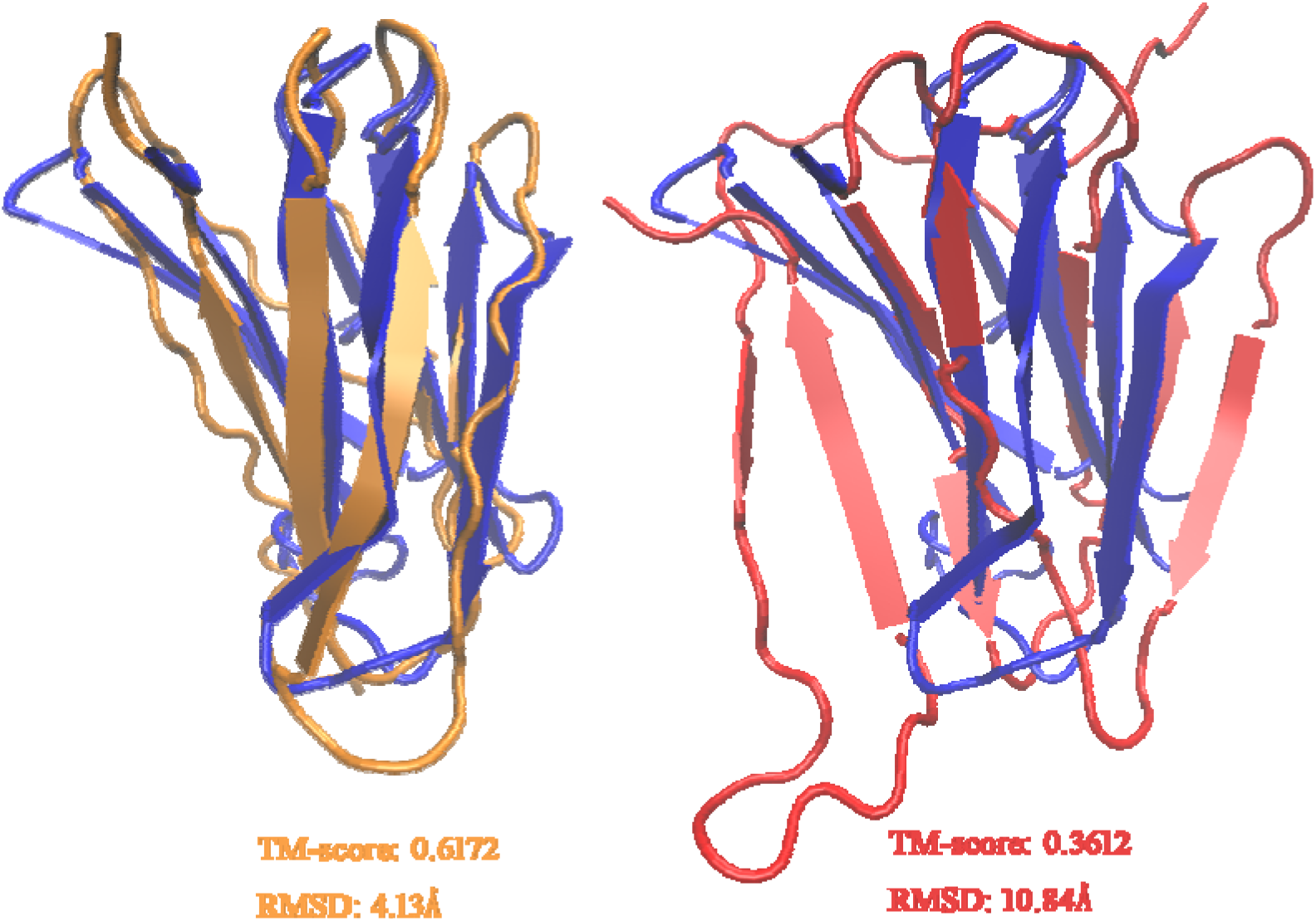
Case study for structure prediction. We illustrate the predicted structures of 1OUSB based on the refined predictions by RDb_2_C (left) and the raw RaptorX-Contact predictions (right), respectively. Comparing to the native structure (blue), the predicted structure based on RDb_2_C (orange) has a higher TM-score (0.6172 *vs.* 0.3612) and smaller RMSD (4.13Å *vs.* 10.84Å) than the predicted structure based on the raw RaptorX-Contact prediction (red).

### Runtime and memory consumption

We evaluated the running time of RDb_2_C on a Dell 5810 workstation (Intel Xeon E5-1620 v3 3.50□GHz CPU, 4 cores, 8 threads and 32 GB RAM) with 8 threads, based on the BetaSheet916 set. Time consumption increases with the size of target protein in a quadratic manner (Fig 10). A typical 400-residue protein needs 20 seconds to complete the prediction. The general memory usage is about 6.3GB. Generally speaking, the runtime and memory usage of RDb_2_C are acceptable for practical protein structure prediction.

**Fig 10.**
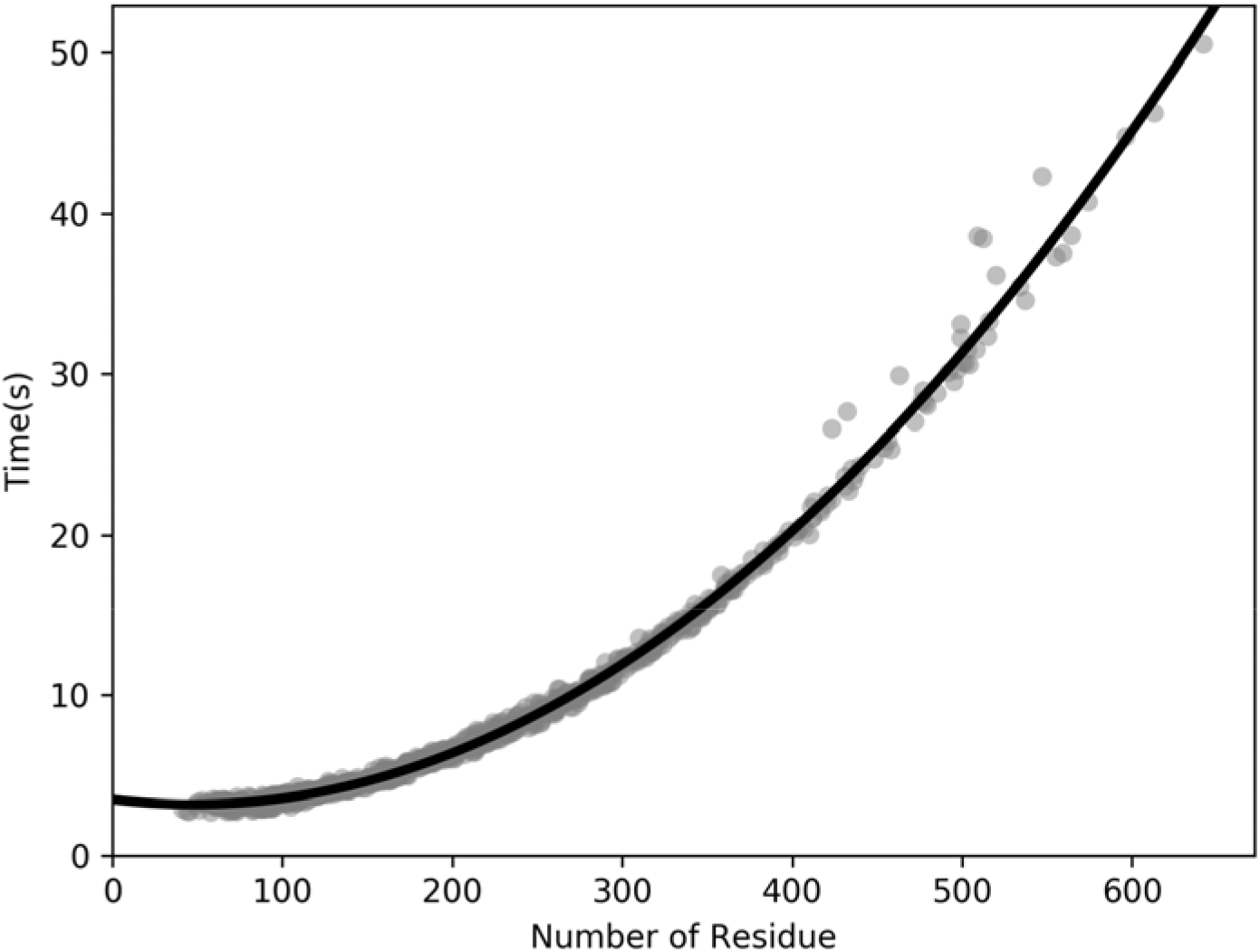
The relationship between runtime and the number of residues. The time consumed increases steadily with the rise of the number of residues (the I/O time is not included).

In conclusion, we developed a ridge-detection-based algorithm with a multi-stage random forest framework to refine the prediction of β-β contacts from a predicted residue contact map. The ridge information could effectively capture the pattern of consecutive residue contacts in interacting β strands. Our method could be pipelined with any residue contact predictors. Tests on CCMpred and RaptorX-Contact suggest that RDb_2_C could improve the prediction of β-β contacts for residue contact predictors of various levels of accuracy. Furthermore, improvement in β-β contact prediction could facilitate the structural prediction of mainly β proteins. The runtime and memory of our method are acceptable for practical use.

## Materials and Methods

### Dataset

We used two well-established datasets for testing: BetaSheet916 [26] and BetaSheet1452 [29]. These two datasets have been widely accepted, thus allowing performance comparison to previous methods. Both datasets were filtered for redundancy. The β residues were defined using DSSP [36], and both β-bridge and extended β-strand residues (B and E in DSSP) were considered as β residues.

We built our training set from the CATH database of protein domain, version 4.1 [39]. Since our work focused on contacts in β strands, only β and α/β domains were considered. In order to eliminate the redundancy between the training set and test sets, we removed all domains from the training set that belongs to the same CATH fold groups as proteins in the two test sets. The fragmented and overly short (< 30 residues) domains were also discarded. Finally, only domains in the CATH S35 set [40] (a subset of CATH with pairwise sequence identity <35%) were kept to reduce the redundancy inside the training set. Thus, there were 493 domains in our training set (Table 10 and S1 Table).

**Table 10.** General information of the training and test sets.

| Numbers | Training set | BetaSheet916 | BetaSheet1452 |
| --- | --- | --- | --- |
| Proteins | 493 | 916 | 1452 |
| Residues | 73580 | 187516 | 361668 |
| $\beta$ residues | 22283 | 48996 | 88352 |
| β-β contact residue pairs | 13278 | 31638 | 56552 |
| $\beta$ strands | 4633 | 10745 | 19186 |
| $\beta$ strand pairs | 2678 | 8172 | 14241 |

In the training set, true β contacts were calculated following the DSSP definition with isolated β-bridge pairs ignored. The DSSP assignment was simplified into 3 categories: H, E and C. The secondary structure probabilities were predicted by DeepCNF [35]. The MSAs were built by HHblits [41] against the UniProt20 database [42], from which residue contact maps were then predicted by CCMpred. ProDy [43] was adopted as a package in Python for dealing with PDB files and analyzing protein structures.

### Ridge Features

We employed the ridge as a proxy to capture consecutively distributed regions of relatively strong signals. The ridge is an extended concept of a local maximum. In an N dimensional space, a local maximum point should be maximal in all N dimensions, while a ridge describes a continuous curve each point of which is the local maximum in the N-1 dimensional subspace orthogonal to the curve. Fig 11A demonstrates a ridge on a 2D image, where the vertical axis stands for the signal strength. Ridge is a good measure to characterize the central axis of an elongated object, i.e. consecutive residue contacts in interacting β strands on a residue contact map.

**Fig 11.**
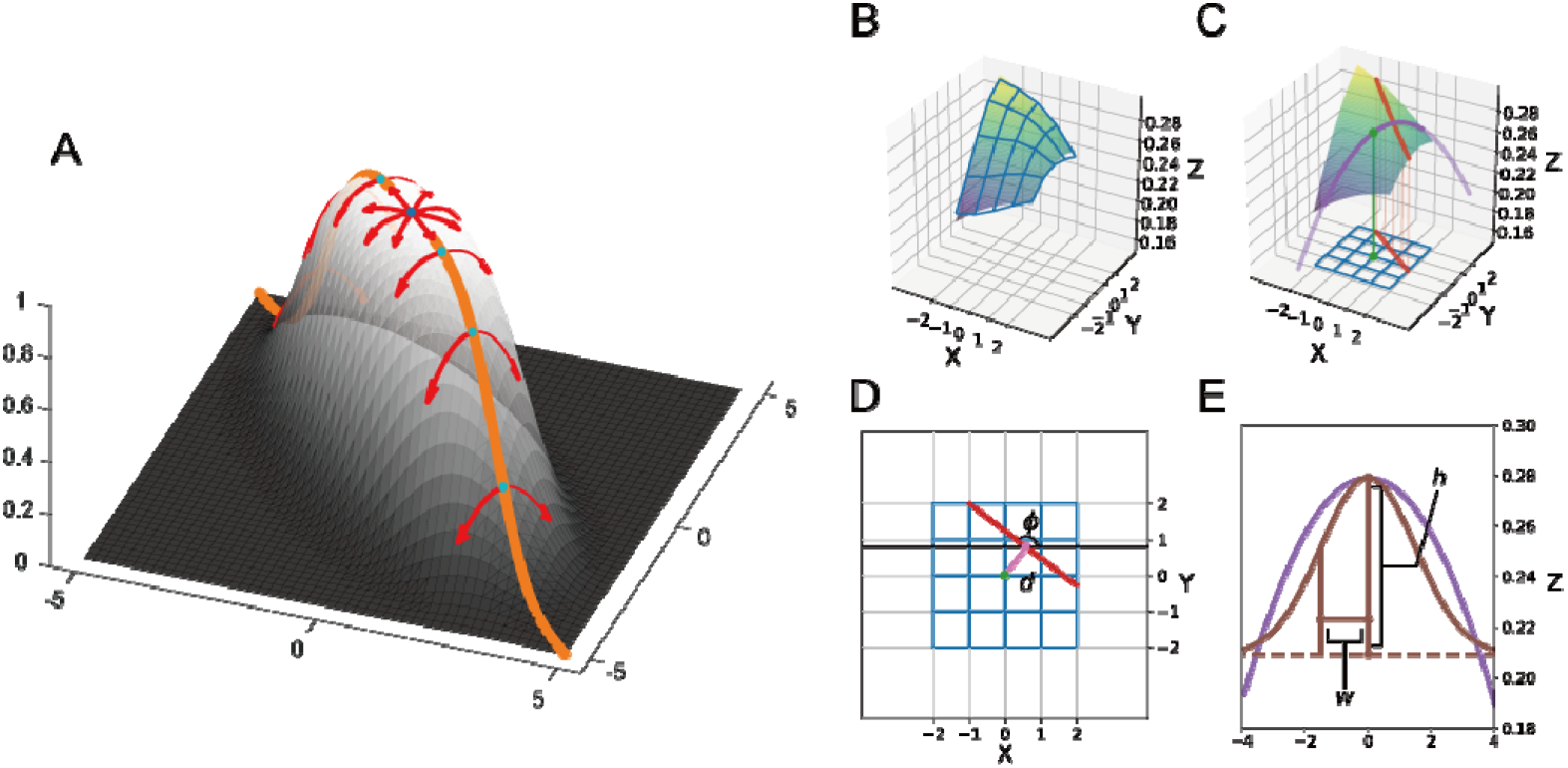
Ridge features from the original map. **(A)** The orange line indicates the ridge on the 2D function surface. All ridge points on the ridge line are the maxima in the directions perpendicular to the line (red arrows). The local maximum point (dark blue) is also a ridge point based on the definition. **(B)** For each given point on the contact map, we select local region (i.e. the grid points) to approximate a quadratic function. **(C)** On the quadratic function surface, we could identify the linear ridge and project it to the XY plane. **(D)** Direction of the ridge □ and distance from the original given point to the ridge *d* could be obtained from the projection. **(E)** We could also identify the principal curvature direction on the ridge and approximate the cross section curve with a Gaussian ridge. The height *h* and width *w* are defined as the height and the standard deviation of the Gaussian function. Details are given in the S1 Text.

For any given point on the 2D map, we firstly estimated the local 1^st^ order and 2^nd^ order derivatives to build the local gradient **∇f** and the Hessian matrix **H** via an ordinary least squares on the extended surrounding region with the size of 5×5. Then we calculated the two principal curvatures (*λ*_*p*_, *λ*_*q*_) by performing eigendecomposition to the Hessian matrix:

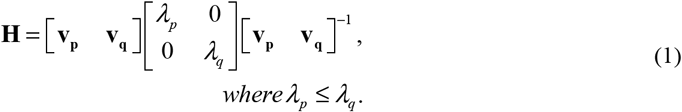

We required at least one principal curvature is negative (i.e. concave) and the directional derivative along the corresponding direction is zero to guarantee the property of ridge points:

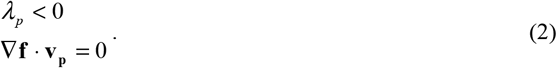

By locating such points on the contact map, we could identify the axis of the elongated region with relatively strong signals.

However, straightforward ridge detection described as above is not practical on discrete maps for several reasons. Firstly, the ridge could not always locate exactly on a discrete point. Secondly, straightforward method will include all ridges without considering the ridge height or strength. For the first issue, we could roughly locate the ridge position by approximating the neighboring region with a quadratic function according to the estimated gradient and Hessian matrix (Fig 11B). Under the approximation, the ridge is a straight line (Fig 11C), from which we could identify the direction (□) and the distance from the original given point (*d*) in the XY plane (Fig 11D). To solve the second issue, we introduced the γ-normalized scale method developed by Lindeberg [34]. In specific, we utilized the square principal curvature difference (*NL*), a measure introduced in Lindeberg’s work, to quantify the ridge strength:

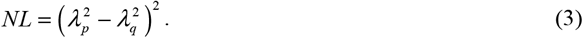

Here, we describe the procedure briefly. We smoothed the map with a Gaussian filter at a series of scale *σ.* However, *NL* is not guaranteed to reach maxima at the scale of the ridge width. Lindeberg introduced γ-normalized *NL* to solve this problem. By multiplying *σ*^*λ*^ with a carefully-selected *γ*, the γ-normalized *NL* could reach maxima at desired ridge width:

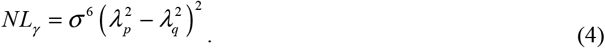

The *γ*-normalized scale method could provide an unbiased estimate of the ridge width (*w*). We further estimate the ridge height (*h*) via a similar process (Fig 11E). More details of the *γ*-normalized scale method and the corresponding calculation protocol in processing contact maps could be found in the S1 Text.

### Model Features

For a given point on the predicted residue contact map, we calculated the ridge features (including ridge direction □, distance to the ridge *d,* ridge height *h* and ridge width *w* (see Fig 11B-E)). These features and scores of the input map jointly constitute 5 N×N matrices (Fig 12). We also incorporated the predicted secondary structure probabilities (for H, E and C) from DeepCNF. Furthermore, to describe positions of the target residue pair, we included the difference in indices of the two residues as well as distances of each residue to both ends of the protein in the amino acid sequence as position features. To characterize the quality of the original contact map, we employed the number of homologous sequences in the MSA per residue as well as the standard deviation of prediction scores as map features.

**Fig 12.**
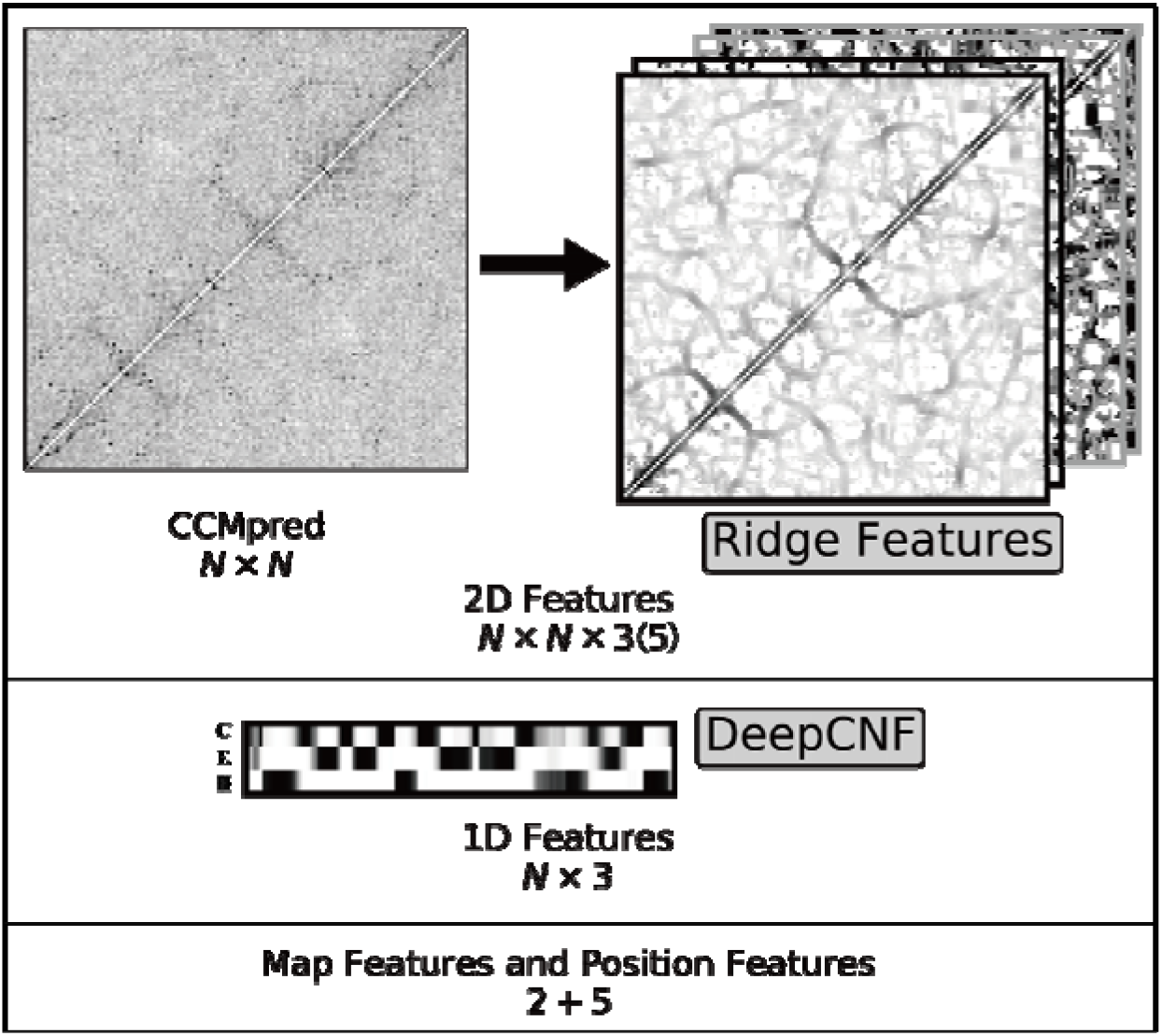
Summary of features adopted in our model. For each target protein with N residues, we have the original CCMpred map with the size of N×N. We calculate the ridge features for each point on the map to get 4 N×N matrices (2 N×N matrices after feature selection). In total, we have N×N×5 (N×N×3 after feature selection) 2D features. The secondary structure prediction from DeepCNF provides an N×3 1D feature matrix. In addition, we have 2 map features (the sequence/residue ratio and CCMpred standard deviation) and 5 position features (1 residue index difference and 4 distances to protein ends). The data in this figure were generated from the protein 1AHQA.

### Model Training and Feature Selection

We applied a 3-stage random forest framework to predict the β-β residue contacts using all features described previously. All random forest models in all stages were set up with 500 decision trees and were optimized by 5-fold cross-validation using the scikit-learn package [44]. The cross-validation was applied in a protein-wise manner, by which the training set proteins were randomly partitioned into 5 mutually exclusive subsets with roughly the same size. Combinations of four subsets were then iteratively used to train the model and to predict the unselected subset. Since all proteins in the training set were predicted independently, the suggested cutoffs were optimized in the cross-validation. Finally, the whole training set was utilized to train a separate model as the final model for evaluation in the test sets.

At the first stage, in addition to features of the target residue pair, we adopted an adjustable window to consider the effect of neighboring residues. Specifically, 2D features (ridge features and the original contact map) of all residue pairs falling within the square window centered at the focus point were included. Secondary structure features of all residues falling within the 1D windows centered at the two target residues were also extracted. Map property features and position features were extracted for the target residue pair only, because they were invariant for the target and neighboring residue pairs. We employed various values of the window size (*ws*), including 1, 3, 5, 7 and 9, to train multiple random forest models at the first stage. Because of the scarcity of β-β residue contacts, the negative (Neg) samples greatly outnumbered the positive (Pos) ones with a Pos/Neg ratio of about 1:600. To simplify the model training, we under-sampled negative samples at different Pos/Neg ratios from 1:1 to 1:40. The under-sampling was implemented in a protein-wise manner. That is, for each protein, the number of negative samples was specifically set based on the number of positive samples. Based on the cross-validation results (Table 11), improvement in model performance becomes saturated at Pos/Neg ratios of 1:40. Therefore, each random forest model was trained at 1:40 Pos/Neg ratios. At the same time, we noticed that the model with the window size of 1 significantly underperforms models of the other window sizes. Therefore, we selected window sizes of 3, 5, 7 and 9 for further model optimization.

**Table 11.** The cross-validation F1-scores for different window sizes and Pos/Neg ratios.

| Pos/Neg | 1×1 | 3×3 | 5×5 | 7×7 | 9×9 |
| --- | --- | --- | --- | --- | --- |
| 1:1 | 32.55% | 37.41% | 37.61% | 35.93% | 34.96% |
| 1:5 | 35.31% | 41.39% | 41.03% | 39.42% | 38.51% |
| 1:10 | 36.31% | 42.65% | 42.00% | 40.76% | 39.39% |
| 1:20 | 36.90% | 43.41% | 43.22% | 41.92% | 40.60% |
| 1:30 | 37.20% | 44.15% | 43.61% | 42.67% | 41.23% |
| **1:40** | **37.33%** | **44.16%** | **44.02%** | **42.90%** | **41.71%** |

Winner in each category is highlighted in bold. The row of the selected Pos/Neg ratio is shown in shadow.

We performed the feature selection by removing features group by group and re-conducting the 5-fold cross-validation. We found that the ridge width *w* and the distance from the ridge *d* are not essential for the model. After removing these two sets of features, only the ridge height *h* and the direction of the ridge □ were kept as ridge features. Thus, we obtained the optimized feature combination as indicated in Table 2. We further optimized the shape of the window. Because **β**-**β** pattern depends on the signals on diagonal and off-diagonal directions, we used the cross-shaped masks with different diagonal width *(dw)* besides the square window mask for 2D features (Fig 13). For all window sizes, the best masks were the ones with the diagonal width of 3 (Table 12). Eventually, we chose the models with the diagonal width of 3 as the final ones.

**Fig 13.**
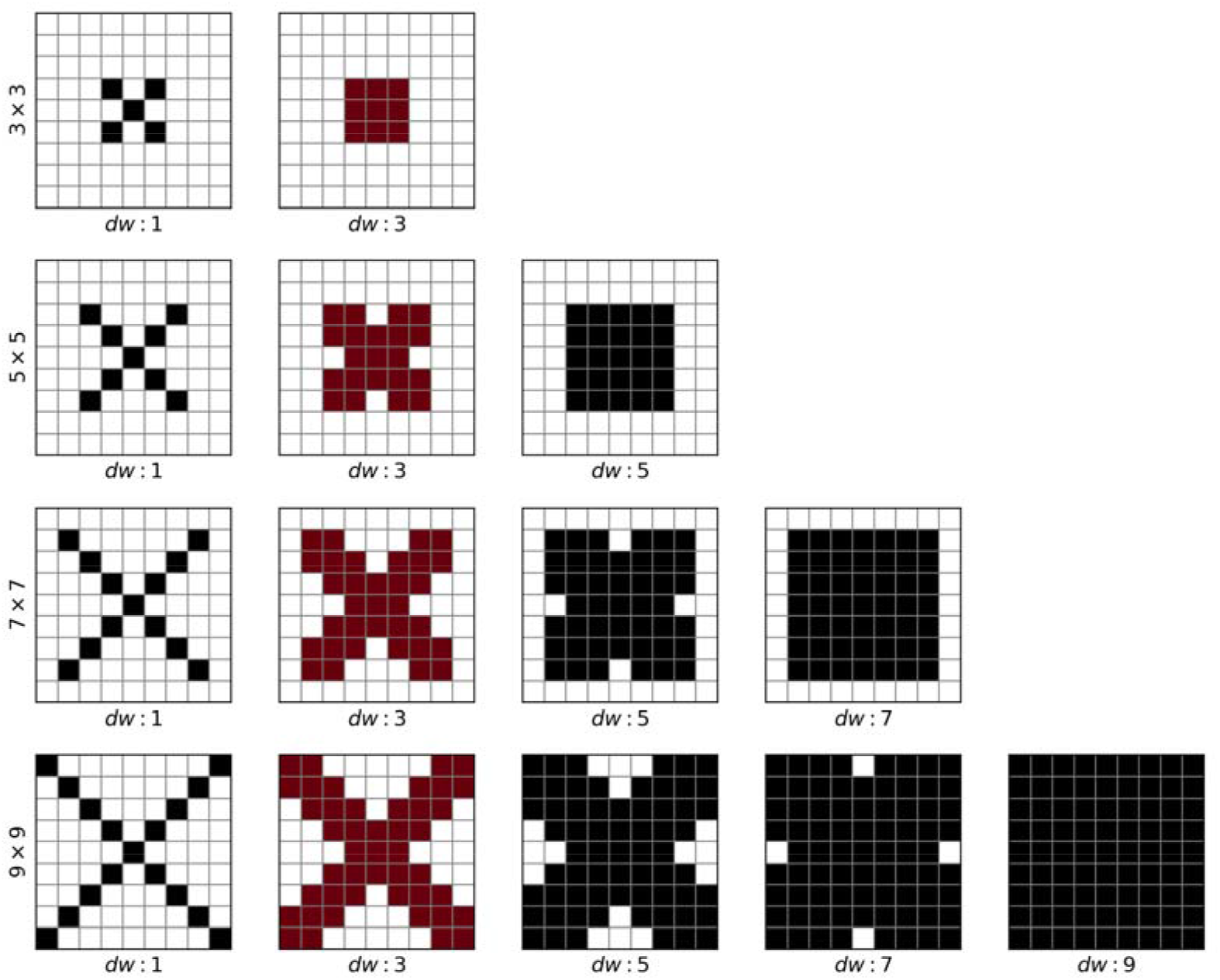
An illustration of the window mask. The selected features are labeled in dark colors. The final window masks that were selected are marked in red.

**Table 12.** The cross-validation F1-scores for different window sizes and diagonal widths.

| Window size | Diagonal width |  |  |  |  |
| --- | --- | --- | --- | --- | --- |
|  | 1 | 3 | 5 | 7 | 9 |
| 3×3 | 41.20% | **44.40%** |  |  |  |
| 5×5 | 41.94% | **45.44%** | 45.27% |  |  |
| 7×7 | 41.12% | **44.80%** | 44.58% | 44.64% |  |
| 9×9 | 40.32% | **44.30%** | 43.95% | 43.55% | 43.36% |

Winner for each window size is highlighted in bold. The column of the selected diagonal width is shown in shadow.

Predictions from the first-stage models were then fed to models at the second stage. In specific, we retained the output scores of the first-stage models as additional 2D features. Unlike the strong constraints applied by bbcontacts that artificially restricts each residue to form no more than two β-β contacts, we included the ranks of each point among the output scores of each column and row and allowed the random forest model to automatically learn the geometry constraints. Hence, output map from each first-stage model provided N×N×3 features (1 N×N raw output and 2 N×N rankings). Subsequently, we performed the feature selection again as the first stage. The first-stage raw scores, the first-stage rankings, ridge features (ridge height *h* and ridge direction □) and predicted secondary structure information by DeepCNF were finally retained after feature selection (Fig 14). The window size and the diagonal width were both optimized at 3 (3×3 square). Then, we combined features from the 4 first-stage models of various window sizes to construct a comprehensive second-stage random forest model. At the third stage, we carried out a similar protocol as the second stage and obtained a final third-stage random forest model.

**Fig 14.**
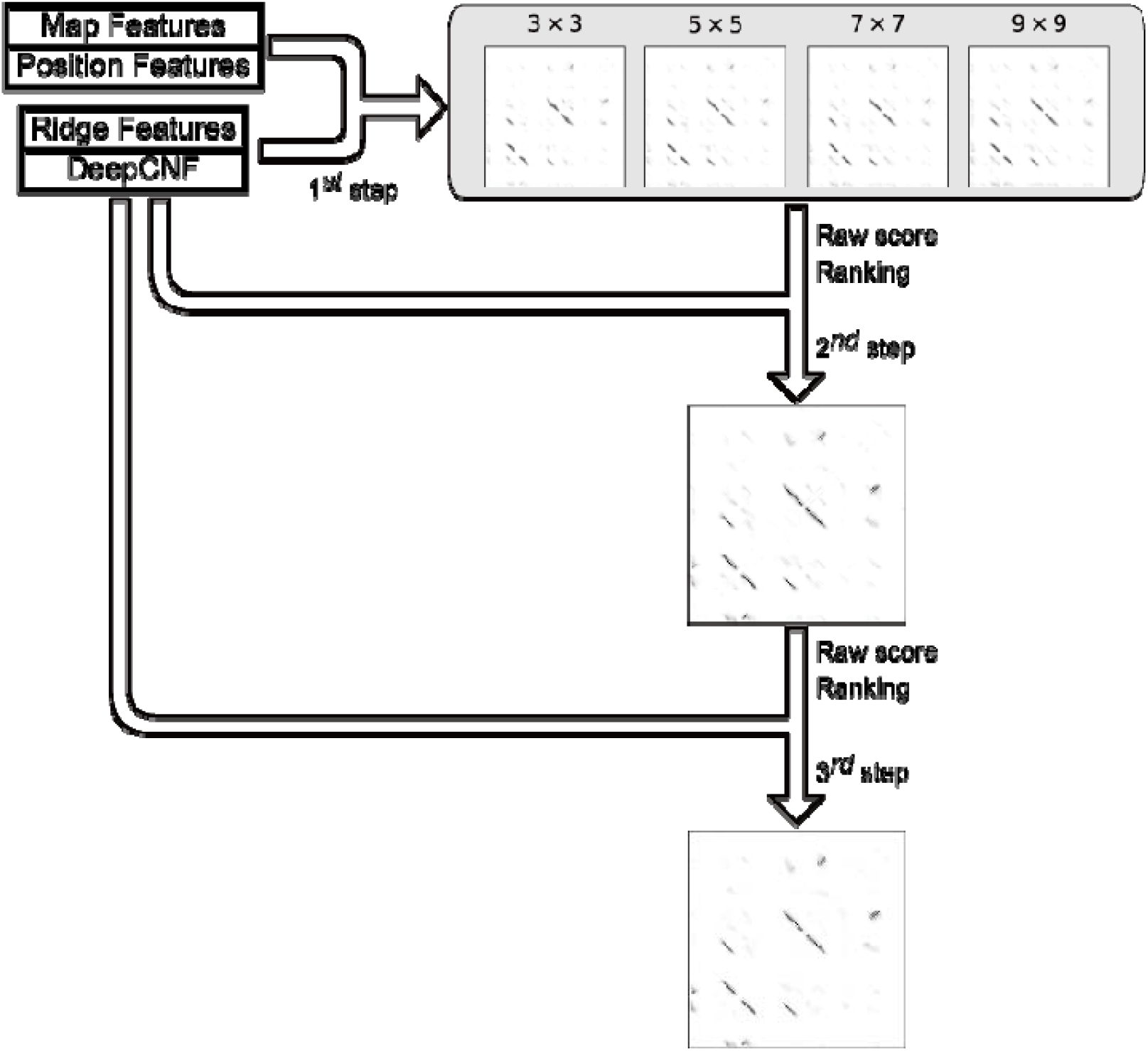
An illustration of the multi-stage framework. In our 3-stage framework, we firstly construct models with different window sizes. We then integrate four models to get the second-stage results. The final result is obtained from the third-stage model. The data in this figure were generated from the protein 1AHQA.

The overall framework was constructed for two different types of secondary structure information, prediction from DeepCNF and assignment from DSSP, respectively. For DSSP-based models, the secondary structure probability is set to 1 for the native category and 0 for the others.

### Evaluation

Performance was evaluated at both residue and strand levels, using measures including Precision, Recall as well as F1-score. Precision and Recall quantify proportions of true positives within all predicted and all native β-β contacts, respectively, while F1-score is the harmonic mean of Precision and Recall:

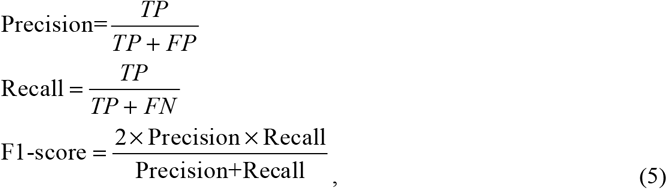

where *TP*, *FP* and *FN* denote true positives, false positives and false negatives, respectively.

Although our method was developed with predicted secondary structure information for practical protein structure prediction, we performed evaluation for models fed with predicted and DSSP-assigned secondary structures respectively to simplify comparison with previous methods. Since bbcontacts is the best method so far and exhibits significantly superior performance to all previous ones, we mainly compared our method with bbcontacts. Results of bbcontacts were obtained following the protocol of the original paper, with secondary structures predicted by PSIPRED [45]. The residue-level evaluation is straightforward, while the strand-level evaluation, however, could only be conducted with the knowledge of clearly defined secondary structures. Thus, we only provide the strand-level results for DSSP-based models. As for the definition of strand pairing, we regard a pair of β strands as interacting if at least one pair of residues on the two strands is predicted as contacting.

### Structure modeling using predicted contacts

All 61 mainly β proteins (with ≥50% of β residues) were chosen from the shrunk BetaSheet916 set (S2 Table), and tertiary structure models of them were constructed with predicted contacts taken as constraints, using the downloadable programs of Crystallography & NMR System (CNS) [46] suite and CONFOLD package [37]. We retained all β-β contacts predicted by the RDb_2_C model in pipeline with RaptorX-Contact at the suggested cutoff as the highly reliable contact pairs, and then enriched the list of contact pairs to 1*L* by collecting the high-ranked and non-redundant RaptorX-Contact predictions from the region outside the predicted β-β region of RDb_2_C (All contacts falling within the square window covering the RDb_2_C prediction points or lines are considered as redundant). These top 1*L* residue contacts were used as distance restraints to fold the protein following the standard CONFOLD protocol, with the DeepCNF results supplemented as predicted secondary structures [37]. A strict restraint of 3.5-6Å was applied to constrain the C_β_ atoms for the more reliable contact pairs of RDb_2_C prediction, whereas a loose restraint of 3.5-10Å were adopted for the non-redundant contact pairs enriched from RaptorX-Contact because these complement pairs are of lower confidence levels. In the control experiment, the top 1*L* residue contacts were directly chosen from the RaptorX-Contact results and a uniform standard restraint of 3.5-8Å was engaged to constrain all contact pairs. For each tested protein, 20 models were generated by CONFOLD, and the 5 models that fit the restraints best were retained. The model with the highest TM-score among the top 5 models was then taken as the representative one for evaluation.

## Acknowledgements

We gratefully thank Prof. Jinbo Xu for his help in the job submission using the RaptorX-Contact server.

## Supplementary materials

**S1 Text. Technical details of the** γ**-normalized scale method for ridge detection and the corresponding calculation protocol in processing contact maps.**

**S1 Table. List of domains in the training set.**

**S2 Table. Results of structure prediction for 61 mainly** β **proteins.**

